# Vibration’s frequency and intensity for optimal setup for enhancement bone response in small rodents: A systematic review and Bayesian network meta-analysis

**DOI:** 10.64898/2026.07.08.737040

**Authors:** Nilson R. S. Silva, Terrell Engmann, Karine J. V. Stoelben, Nurbanu Bursa, Alex X. Zhang, Kurt S. Soloniuk, William R. Thompson, Jung Min Hong, Gunes Uzer

## Abstract

Low-intensity vibration (LIV) is a non-invasive mechanical stimulus capable of regulating skeletal adaptation and cellular signaling pathways involved in bone remodeling. Despite growing interest in LIV, substantial methodological heterogeneity persists in the selection of experimental vibration parameters such as frequency, expressed in Hertz (Hz) and intensity, defined as earth’s gravitational field (g) (9.81 m/s^2^). Focusing on micro-computed tomography (µCT) derived trabecular bone volume fraction (BV/TV) as the main outcome mesure, this study sought to synthesize the effects of different LIV frequency and intensity on BV/TV in small rodents (mice and rats) as they remain as the most studied pre-clinical model. To accomplish this, we performed a systematic review searching for publications in English on PubMed, Web of Science, CINAHL, and Embase databases. Two independent investigators followed inclusion criteria to select only peer-reviewed studies with mature mice, using whole-body vibration experiments without other co-variables. We further restricted to include studies that analyzed non-fractured bones and compared pre- and post-intervention or control values. In addition to these core criteria, a detailed hierarchical screening framework was applied during full-text review. The two independent investigators extracted data independently and considered the characteristics of the study, animals’ characteristics, intervention characteristics, and results. For this study we considered load-bearing hindlims, femur and tibia, separately but did not include vertebrae in the analysis. A Bayesian network meta-analysis and a revised SYRCLE risk of bias (RoB) tool were used to evaluate the risk of bias across included studies. Seven studies met the inclusion criteria. Results showed that an LIV regime applied at 45Hz at 2g presented higher chances to increase trabecular BV/TV of the mouse tibia (estimated effect 3.22 [CrI 1.98, 4.45]), while LIV regimes applied to the femur at 90Hz and 1.4g (estimated effect 3.08 [CrI -1.99, 7.97]) present better chances to increase trabecular BV/TV results compared to other interventions but with no significant differences. Finally, we applied 45Hz at 0.2g LIV to 5 month old male C57BL/6 for 5 weeks (n=10/group) which showed significantly increased Trabecular Thickness (Tb.Th) for both the tibia (10%, p<0.01) and femur (17%, p<0.001), with the femur showing further increaseses in trabecular BV/TV (32%, p<0.05) compared to non-LIV controls. We conclude that changes in the microarchitectures of the tibia and femur respond differently to the same application of LIV (45Hz, 0.2g) in mice and rats.

## INTRODUCTION

As a non-invasive and easy-to-apply strategy, Low-intensity vibration (LIV) has gained significant research attention as a non-pharmacological intervention for enhancing and maintaining musculoskeletal; competence^1^. LIV is a low impact, high frequency (30-500Hz) mechanical regime originaly developed to emulate the power spectrum of muscle contractility during exercise^2^. In contrast to exercise, however, which can induce peak strain of 3,500 microstrain(µε)^3^ and 200 µm/s peak fluid velocities^4^ in bone, LIV-induces matrix deformation as low as 50µε^5^. As such, LIV may exert its effects through various other mechanisms.

First, dynamic accelerations caused by LIV generates fluid shear stress on bone surfaces that can reach up to 2Pascals (Pa) due to high viscosity of the bone marrow^6–8^. Fluid shear stress is thought to be one of the primary factors governing alterations in trabecular bone mechanical force^9,10^. Second, LIV promotes muscle contraction, which in turn generates tension on the periosteum and thus the bone surface^11–15^. Finally, emerging evidence from our lab^16^ and others^17^ shows that LIV directly-induces motion of the nucleus relative to cell body. While the exact mechanism(s) though which LIV influences cellular function remains a topic of research, two recent meta-analyses of clinical trials concluded that LIV increases bone density in both healthy and post-menopausal women^18,19^. Other randomized clinical trials show that populations unable to engage in rigorous physical activity—including patients with Anorexia Nervosa^20^, cancer^21,22^, and Duchenne Muscular Dystrophy^23^ all benefit from LIV with improvements in bone quality. While LIV has shown promising results, there remains a lack of consensus regarding the experimental parameters used in LIV namely the frequency at which LIV is applied and the acceleration magnitude, usually defined in g where 1g is the Earth gravitational field (9.81 m/s^2^). Owing to large variation of protocols for *in vitro* and clinical studies, here we sought to understand the most effective frequency/acceleration combinations in pre-clinical mouse models as they remain the most largely adopted model with relatively uniform experimental design^24,25^. Previous small rodent studies studies have employed a range of frequencies (e.g., 30Hz, 32Hz, 45Hz, 90Hz) and intensities (e.g., 0.10g, 0.15g, 0.30g)^26–31^, yielding inconsistent results^5,22^. This methodological heterogeneity in LIV’s frequencies, intensities, and outcomes highlights how comparisons across studies and conclusions are difficult to generalize. Establishing standardized LIV parameters in small rodent models would help improving the reliability of LIV approaches. A variety of commonly used techniques, such as µCT, histology, and mechanical testing, are used in preclinical rodents models to evaluate bone response^32,33^. Among the available µCT parameters, the ratio of bone volume to tissue volume (BV/TV) is widely used in bone microarchitecture investigations^34,35^. We chose to compare studies using BV/TV as it not only offers a robust and quantifiable reproducible index of the bone volume of interest at a given time point, but is also a standard output available across most µCT systems^32,36^. We also decided to investigate femurs and tibias, as the distal femur and proximal tibia are the most frequently studied sites in small rodent LIV studies, given their abundance of metabolically active trabecular bone and their well-established site-specific responsiveness to mechanical stimuli^27,37–40^. Therefore this study aimed to synthesize the available data to determine the most promising combinations of LIV parameters to improve BV/TV in small rodents^22,23,24^. To this end we conducted a systematic review and Bayesian network meta-analysis to identify which vibration frequency–intensity regime produces the most effective bone response. To account for the heterogeneous evidence across rodent studies, we chose to analyze the bone tissue volume data using a Bayesian ranking framework. Our approach enabled us to integrate the data and generate probabilistic estimates of comparative main effects and credible intervals (CrI). Finally, we performed a study using one of the most promising LIV protocols found in our ranking to evaluate its effects in bone morphonlogy in mice.

## METHODS

### Overview

We focused on investigating how different frequencies and intensities of LIV affect bone tissue. To do so, we divided our investigation into two parts. Initially, we conducted a systematic review and Bayesian network meta-analysis following the Preferred Reporting Items for Systematic Reviews and Meta-Analysis (PRISMA) Statements guidelines^41^. Next, we experimentally compared LIV outcomes for the most and the least probable efficient LIV setups for BV/TV enhancement. The systematic review section of this research was registered in PROSPERO data base (ID: CRD420251000231).

### Information sources, selection process and eligibility criteria

Starting on April 18th, 2025 an independent researcher (NRSS) conducted a comprehensive search on PubMed, Web of Science, EMBASE and CINAHL, using a series of keywords employing PICO strategy (see Supplemental material - Databases Search Strategies). We included full-text studies in English. After removing duplicates, two researchers (NRSS & TE) conducted a comprehensive independent screening process on the selected papers starting by excluding returns based on title and abstract. Once screening was done, the researchers included all studies that at least one investigator agreed on including. Then the two researchers conducted a full reading filtering process using specific eligibility hierarchical criteria (see Supplementary Material – Eligibility criteria). We used the Covidence website for filtering management while employing our eligibility criteria. A third researcher (KJVS) was consulted in cases of disagreements during filtering.

### Data collection process

Once the results were filtered, two investigators (NRSS & TE) extracted relevant data points from each study (Table 2) related to training, rodent strain, and bone morphology outcomes like bone volume ratio. We chose to focus on pre and/or post BV/TV ratio as it is considered a primary measure of bone mass widely used in osteoporosis research^42,43^. In cases where the studies did not explicitly provide the mean and standard deviation, we contacted the authors for the results. When contacting the author was not possible, we utilized ImageJ software to extract these values from graphical representations and figures within the articles. For studies that included multiple LIV training groups, each intervention was analyzed separately. If a study featured both a LIV group and another type of intervention (e.g., conventional strength training), we considered only the data from the control and LIV intervention groups.

**Table 1.**
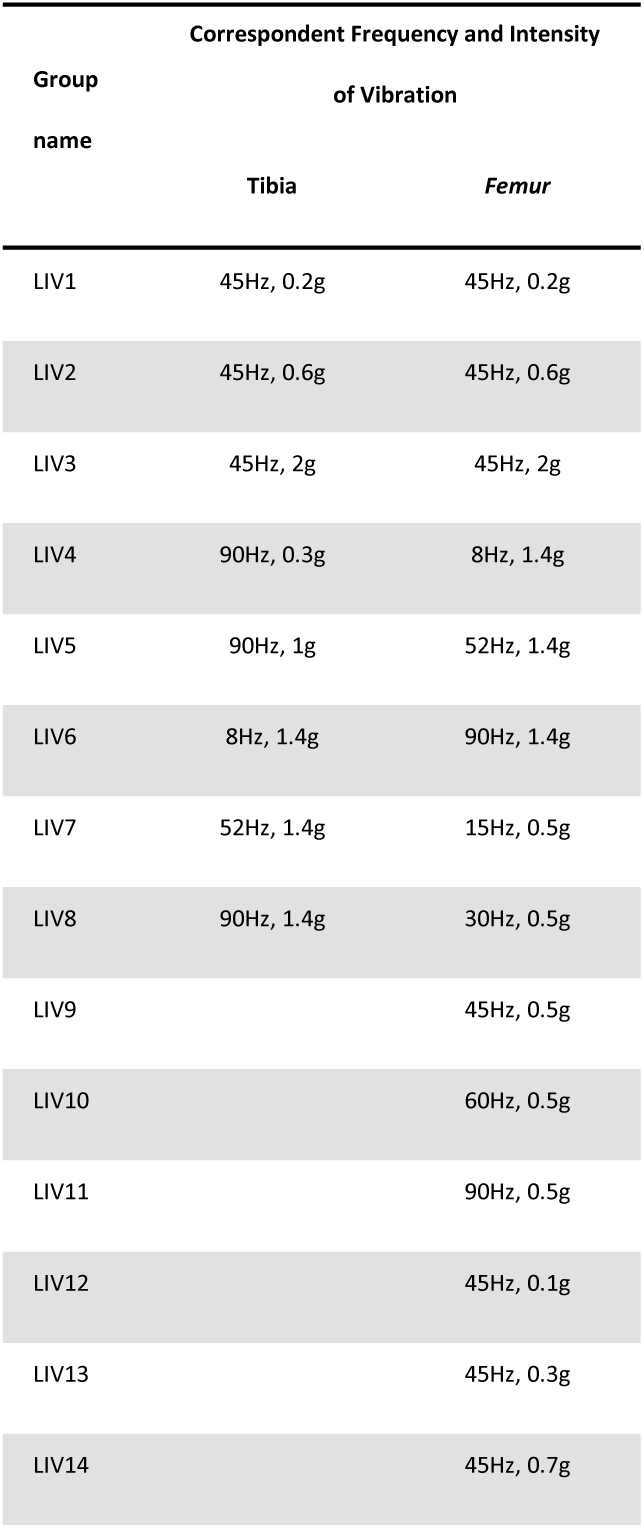
Low intensity vibration regimens selected for analysis for tibia and femur.

**Table 2.**
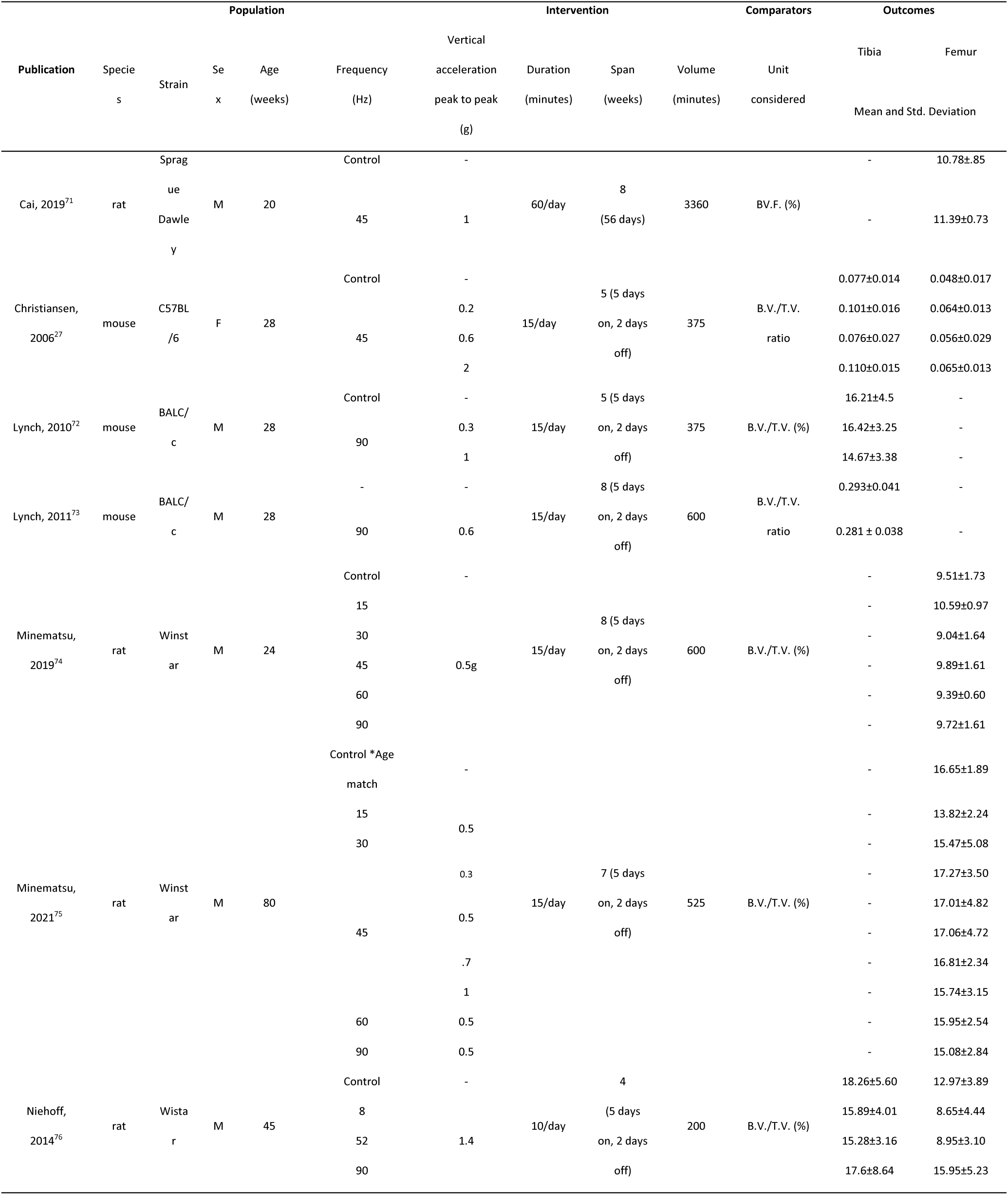
PICO (Population, Intervention, Comparators, Outcomes), collect across select studies.

### Risk of Bias Assessment

For this review, we assessed the risk of bias of the included studies using SYRCLE’s Risk of Bias (RoB) tool for animal studies^44^. This tool is adapted from the Cochrane Collaboration RoB tool which is used in assessing biases in human clinical trials. The screened studies were independently assessed using the ten questionnaire entries identified in the tool which are used to measure six biases including selection bias, performance bias, attrition bias, detection bias, reporting bias, and other biases (Figure 3). Two reviewers (NRSS, TE) independently conducted the risk of bias evaluations to ensure objectivity and reliability. A third researcher (GU) was consulted in cases of non-consensus.

### Bayesian network meta-analysis

Since standard meta-analysis are limited to comparing only two interventions at a time, and some of the included studies evaluated more than two LIV protocols, we opted to conduct a network meta-analysis (NMA)^45^. After comparing fixed and random effects models in the network meta-analysis, we chose to use fixed effects model to work with tibia data, and random effects model to work with femur data, given the results for Deviance Information Criteria of each data set (Supplementary Fig. 1A and 1B). In order to facilitate the random effects NMA, we decided to use a Bayesian statistical approach instead of the frequentist approach, as it handles small sample sizes effectively and provides a straightforward ranking of the different LIV levels through posterior probabilities^46–51^. The fixed and random effect models were adjusted by using three Markov chain Monte Carlo (MCMC) chains and 10,000 iterations for MCMC chains with 1000 iterations burn-in and adaptation periods. The measured stability of the model, as well as the inconsistency assumption, indicates a good convergence range and has shown no violation in all the assumptions, respectively, supporting the validity of the network meta-analysis framework (Supplementary Table 1 A-B; Supplementary Fig. 2A-B, Supplementary Fig. 3A-B). In order to determine the probability of best effectiveness for each LIV treatment, we compared each LIV treatment with an arbitrary common control group (Control) – we calculated the proportion of Markov chain iterations in which each LIV level had the highest probability of being ranked as the most effective, second most effective, and so forth.

### LIV Experiment Design

In order to demonstrate the possible LIV effect over bone microarchitecture, we ran an experimental study using one of the most effective frequencies found by our NMA. We designed the experiment with two groups: LIV at 45Hz, .2g ((+)LIV) and control group ((−)LIV), which were subjected to same conditions with no vibrations. Each of the groups had 10 C57BL/6 male mice, based on an a posteriori power analysis^52^. We subjected the experimental animal groups to 4 weeks (5 days on followed by 2 days off) of LIV for 20 minutes/day. Once the experiments were complete, we euthanized the animals and extracted the hindlimb leg bones (tibias and femors) for micro-CT analysis.

### Micro-CT

We fixed the left femora and tibiae using neutral buffered formalin for 24 hours and then stored emerged in 70% EtoH and kept at 4 degrees for later evaluation. We performed the scan using an X-ray microtomography (Scanco uCT-35) with the following settings: 50 kV, a voxel size of 9 to 13 to μm³ depending on the bone segment. Bone specimens were scanned, reconstructed, and analyzed following previously described methods^53^. We measured parameters for both trabecular and cortical bone^54^.

### Experimental statistical analysis

We used T-Student test to compare the results among groups when parametric assumptions were satisfied - normality was assessed by the Shapiro–Wilk test and homogeneity of variances evaluated using Levene’s test. When assumptions were violated, we used Mann-Whitney U test as a non-parametric alternative.

## RESULTS

### Network Meta-Analysis Identifies Frequency- and Intensity-Dependent Effects of Low-Intensity Vibration on Bone Volume Fraction

Each combination for frequency and intensity included in this study (Table 1) were analysed using a network map (Figure 2 A-B) where each node represents an LIV treatment (LIV1 to LIV14) or the control group (central node) with gray lines indicating direct comparisons between LIV treatments from the included studies. It is worth to highlight that Table 1 should be used as an index when analysing other graphics so you can understand what protocol correspond for LIV# at a given bone. The thickness of the lines connecting the nodes represents the level of similarity between treatments, the thicker the line, the more similar the treatments were. For tibia (Fig. 2C) the forest plot indicated that LIV1 and LIV3 have a estimated positive effects of 2.32 (95% CrI 1.01, 3.62) and 3.22 (95% CrI 1.98, 4.45). While LIV1 and LIV3 values were entirely past to the right of null value - indicating enough effect for increased BV/TV - the other LIV groups presented CrI values overlapping the null value line, meaning not enough effect to grant increased or decreased BV/TV. For femurs, the forest plot (Fig 2D) showed that LIV6, LIV3, LIV1, and LIV13 presented the most positive estimated effects compared to other LIV regimens, with values of 3.08 (95% CrI -1.99,7.97), 1.69 (95% CrI -2.06, 5.32), 1.60 (95% CrI -2.18,5.26), and 1.36 (95% CrI -2.64, 5.33) respectively. Still, according to the the forest plot, none of the LIV groups used for femur presented enought effect for BV/TV to be considered increased or decreased. We determined the probability of each LIV setup to be the most effective by comparing it with an arbitrary common control group. We found such probability by counting the proportion of iterations of the Markov chain in which each LIV level had the probability of being ranked as most effective, second most, and so on. We use surface under the cumulative ranking (SUCRA) scores (Fig. 2D-E) to summarize the relative ranking probabilities of interventions, where higher values indicate a greater likelihood of superior performance. Regarding to tibia, we show once more in Fig 2D that LIV1 and LIV3 outperform the other regimens rankings of probability of performing better. We used such data to generate LIV regimen rankograms (Supplementary Fig 4 A-B) associated with the most or least effective. According to our results tibia’s LIV3 (45Hz, 2g), treatment presented a probability of 90.7% of achieving 1st on ranking, indicating it is likely the most effective treatment followed by LIV1 80.8% (45Hz, 0.2g), while LIV7 (52Hz, 1.4g) with 19.6% is likely to be ranked among the lowest, suggesting it to be the least effective treatment. Meanwhile, the femur data has shown that its LIV 6 (90Hz, 1.4g) had the highest probability (84.4%) of achieving first on ranking, indicating it is likely the most effective treatment, while femur’s LIV4 (8Hz, 1.4g) is likely to be ranked among the lowest (7.6%), suggesting it is the least effective treatment. To summarize the pairwise treatment comparisons across the multiple interventions in terms of effectiveness, we generated a heatmap for the league table (Supplemental Figures 5 and 6), displaying both the upper and lower triangles. Each cell contains the mean difference along with the corresponding 95% credible intervals for the pairwise comparisons between treatments. The columns are arranged from the best to the worst. Taken together, the visual results suggest that for tibial BV/TV, LIV3 (45Hz, 2g) and LIV1 (45Hz, 0.2g) significantly outperform other treatments. On the other hand, the results presented in Figure 2D, 2F and 3B, indicates that across the results involving studies that looked into the LIV effect on femur’s BV/TV, there was only a tendency of some LIV regimens outperform others.

**Figure 1.**
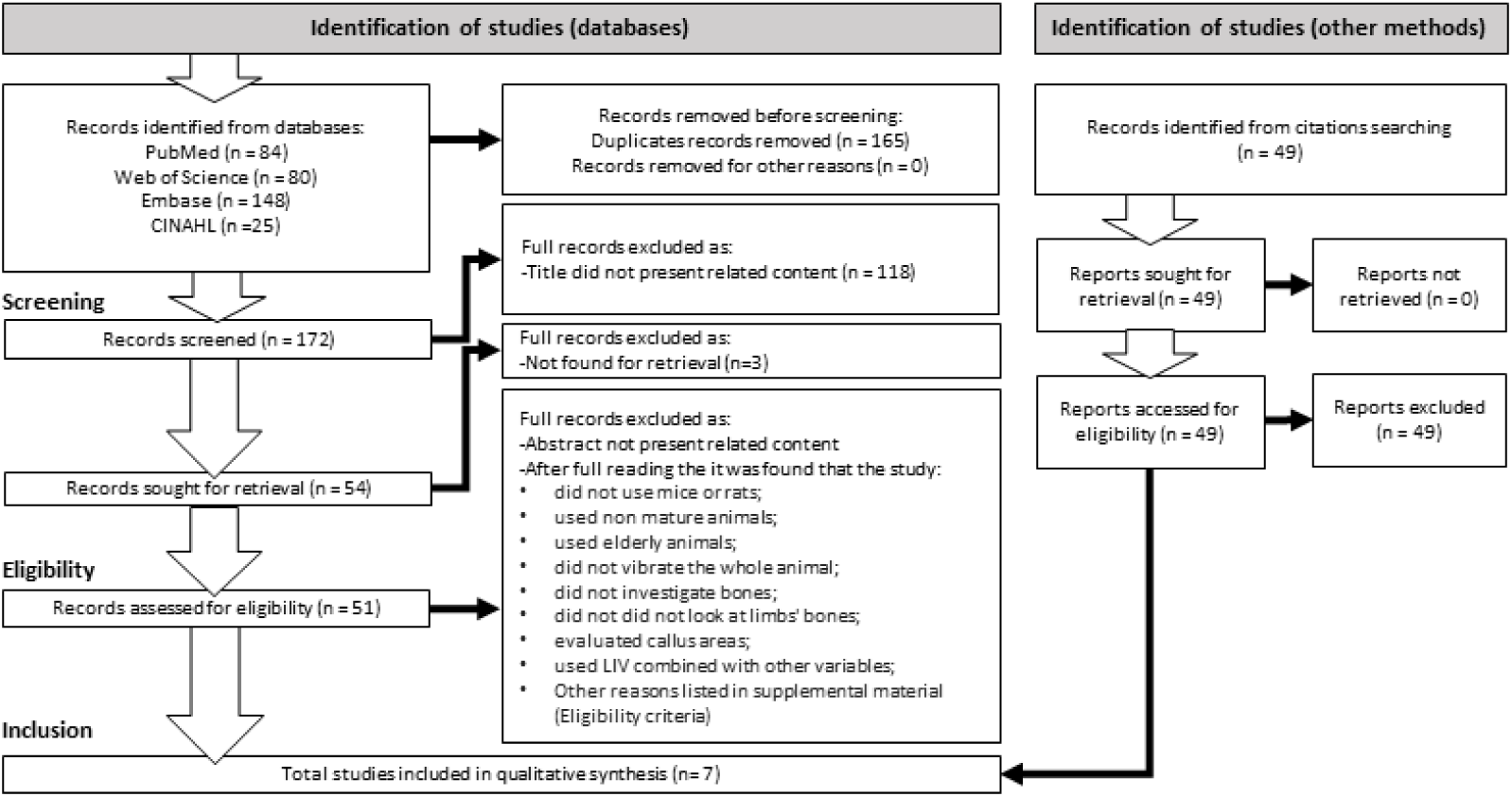
Flowchart for study selection

**Figure 2.**
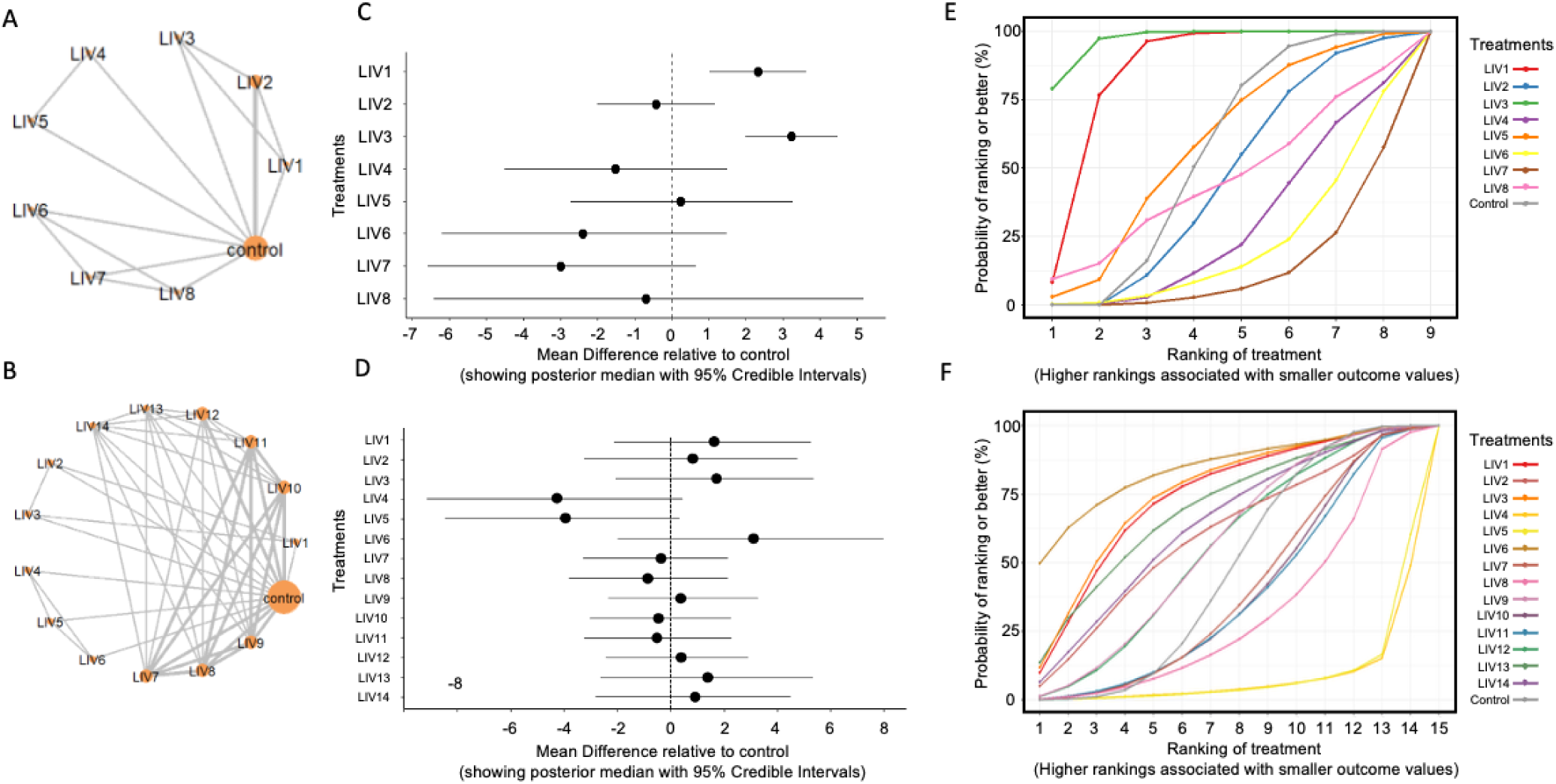
Network and forest and Ranking plots for tibia and femur. (A) Network plot for different LIV regimes on tibia; (B) Network plot for different LIV regimes on femur; (C) Forest plot for different LIV regimes on tibia; (D) Forest plot for different LIV regimes on femur.; (E) Ranking of treatment for different LIV regimes on tibia; (F) Ranking of treatment for different LIV regimes on tibia;

**Figure 3.**
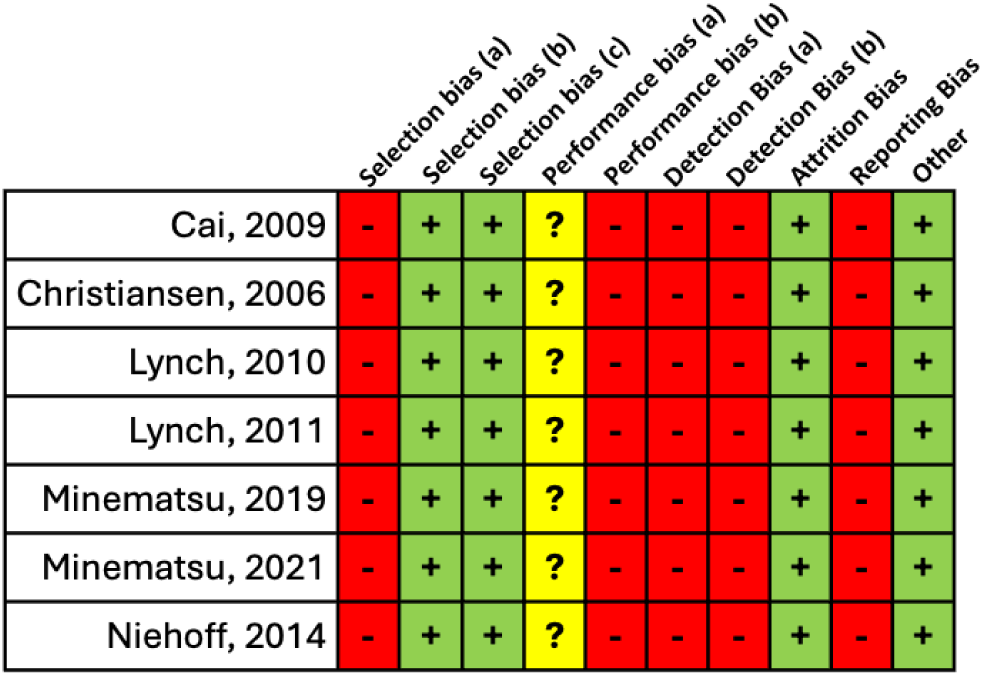
SYRCLE Bias assessment results. Red minus (−): High risk of bias; Yellow question mark (?): Uncertain risk of bias; Green plus sign (+): Low risk of bias. Selection bias (a) Sequency generation, (b) Baseline characteristics, (c) Allocation concealment; Performance Bias (a) Random housing, (b) Bliding; Detection bias (a)Random outcome assessment, (b) Bliding.

### Risk of Bias (RoB) Assessment Reveals Moderate Reporting Limitations Across Included Studies

Following the independent assessment of all metrics presented at SYRCLE RoB tool, the reviewers (NRSS & TE) came to a consensus on each entry, and assigned a bias score of high, low, or uncertain risk of bias. Overall, all the seven include studies scored similarly. Reasons for high risk of bias scores were related to generation and application of an allocation sequence, outlining efforts to blind personnel when necessary, or a lack of access to animal study protocol material. Besides most of these studies are published and indexed on reliable databases, authors failed to provide enough information regarding the details described above. It is important to note that, under SYRCLE RoB tool, a published animal study may only be classified as low bias when explicitly reported certain procedures, even though such level of detail remain relatively uncommon in animal reseach reports. The included papers scored uncertain regarding the randomization of housing. Unfortunately, animal housing details are often insufficiently reported in published animal studies^55^. On the other hand, the studies presented “Low risk of bias” (in green) for “selection bias of baseline characteristics”, “selection bias for allocation concealment”, “attrition of bias” and “Other” which covers studies apparently free of other problems that could result in high risk of bias.

### Low-Intensity Vibration Shows Differential Effects on Femoral and Tibial Bone Microarchitecture

Micro–computed tomography (μCT) demonstrated significant vibration-dependent effects on femoral bone structure (Fig. 5). Vibrated group exhibited a significant increase (+14.65, p<0.01) in total bone mineral content (Tot.BMC, Fig. 4B) as well as (+16.02%, p<0.01) for total tissue volume (Tot.BV, Fig. 4C), and (+17.05%, p<0.01) total bone volume (Tot.TV, Fig. 4D). Trabecular parameters included increased (+14.89%, p<0.01) trabecular thickness (Tb.Th., Fig. 4F), a substantial increased (+62.00%, p<0.01) trabecular bone mineral content (Tb.BMC, FIG. 4I), and greater (+28.57%, p<0.01) trabecular bone volume (Tb.BV/TV, Fig. 4J). Even so, we did not observe significant changes (+4.21%, p=0.188) in trabecular spacing (Tb.Sp, Fig. 4G), and (−4.4%, p=0.116) trabecular number (Tb.N, Fig. 4H), when compared to non-LIV control. For cortical bone, vibrated mice presented a greater (+12.02%, p<0.01) cortical bone volume (Ct.BV, Fig 4K), as well as greater (+17.05, p<0.01) cortical tissue volume (Ct.TV, Fig. 4L). Despite the increases in both bone and tissue volume, cortical bone fraction (Ct.BV/TV, Fig. 4M) presented a small but significant decrease when compared to controls (−4.18, p<0.05). Collectively, our femoral outcomes indicate that the chosen LIV stimulation was associated with significant femur improvements in both trabecular and cortical bone microarchitecture measures compared with non-vibrated animals. For tibial measures, a few variables related to total and cortical bone presented similar results compared to the femurs. We observed increased Tot.BMC (+8.41, p<0.05, Fig. 5B), Tot.BV (+8.16%, p<0.05, Fig. 5C), and Tot.TV (+6.32%, p<0.05, Fig. 5D) when compared to controls. In regards to trabecular related changes (Fig 5E-J), no significant differences were found. With respect to tibia cortical portions, both Ct.BV (+7.67%, p<0.05, Fig. 5K) and Ct.TV (+6.32%, p<0.05, Fig. 5L) presented greater values compared to control.

**Figure 4.**
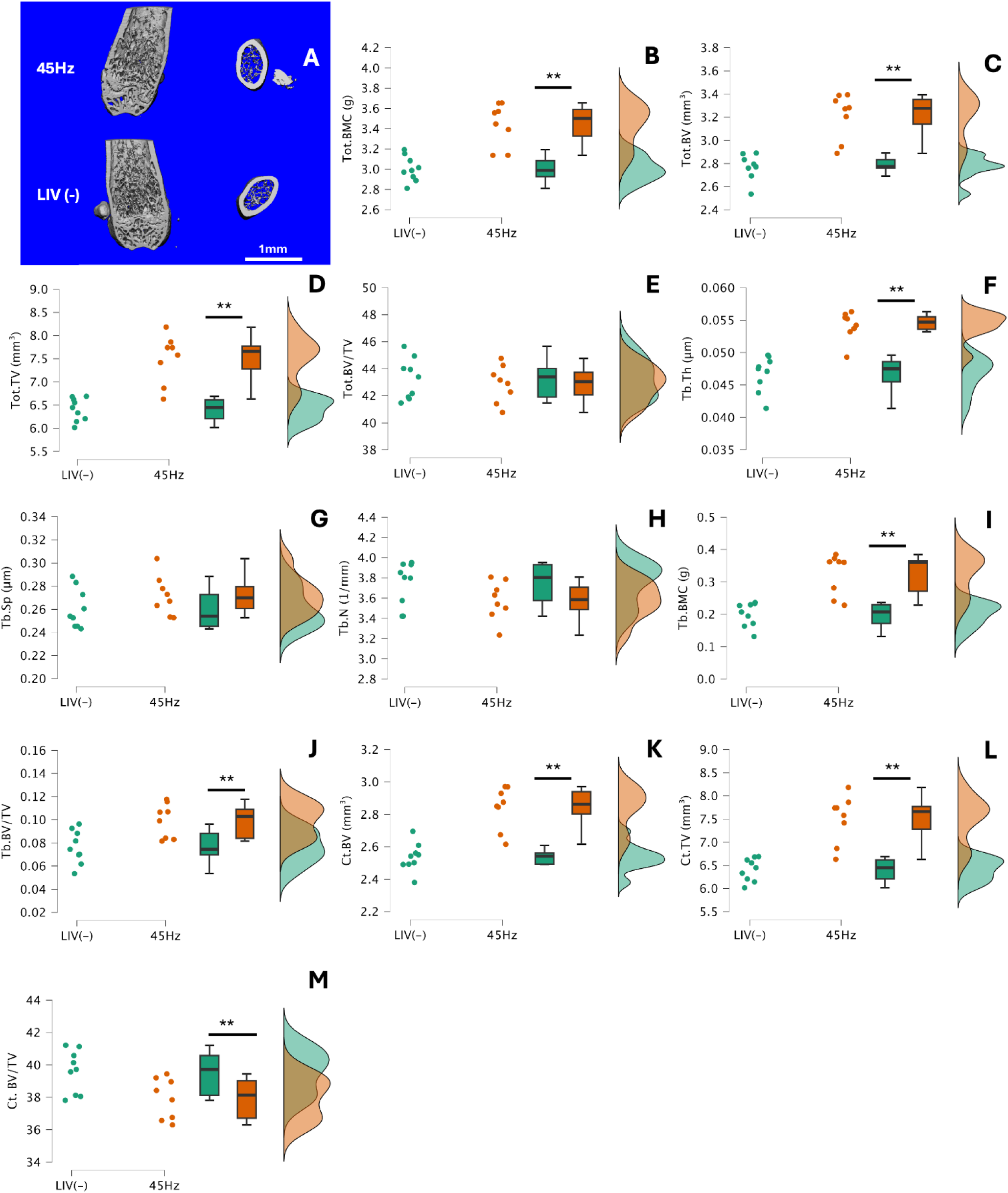
μCT measures derived from femurs of 24weeks old C57BL/6 mice subjected to LIV regime. (A) Representative 3D reconstruction of distal metaphyseal trabecular region of interest (coronal cut) and mid-shaft cortical cross-section of the femur’s region of interest; (B) Total Bone Mineral Content; (C) Total Bone Volume; (D) Total Tissue Volume, (E) Total Bone Volume Fraction; (F) Trabecular Thickness; (G) Trabecular Spacing; (H) Trabecular Number; (I) Trabecular Bone Mineral Content; (J) Trabecular Bone Fraction; (K) Cortical Bone Volume; (L) Cortical Tissue Volume; (M) Cortical Bone Fraction;. Significance between groups tested using T-Student test or Mann-Whitney-U test when not parametric, *p< 0.05, ** p< 0.01

**Figure 5.**
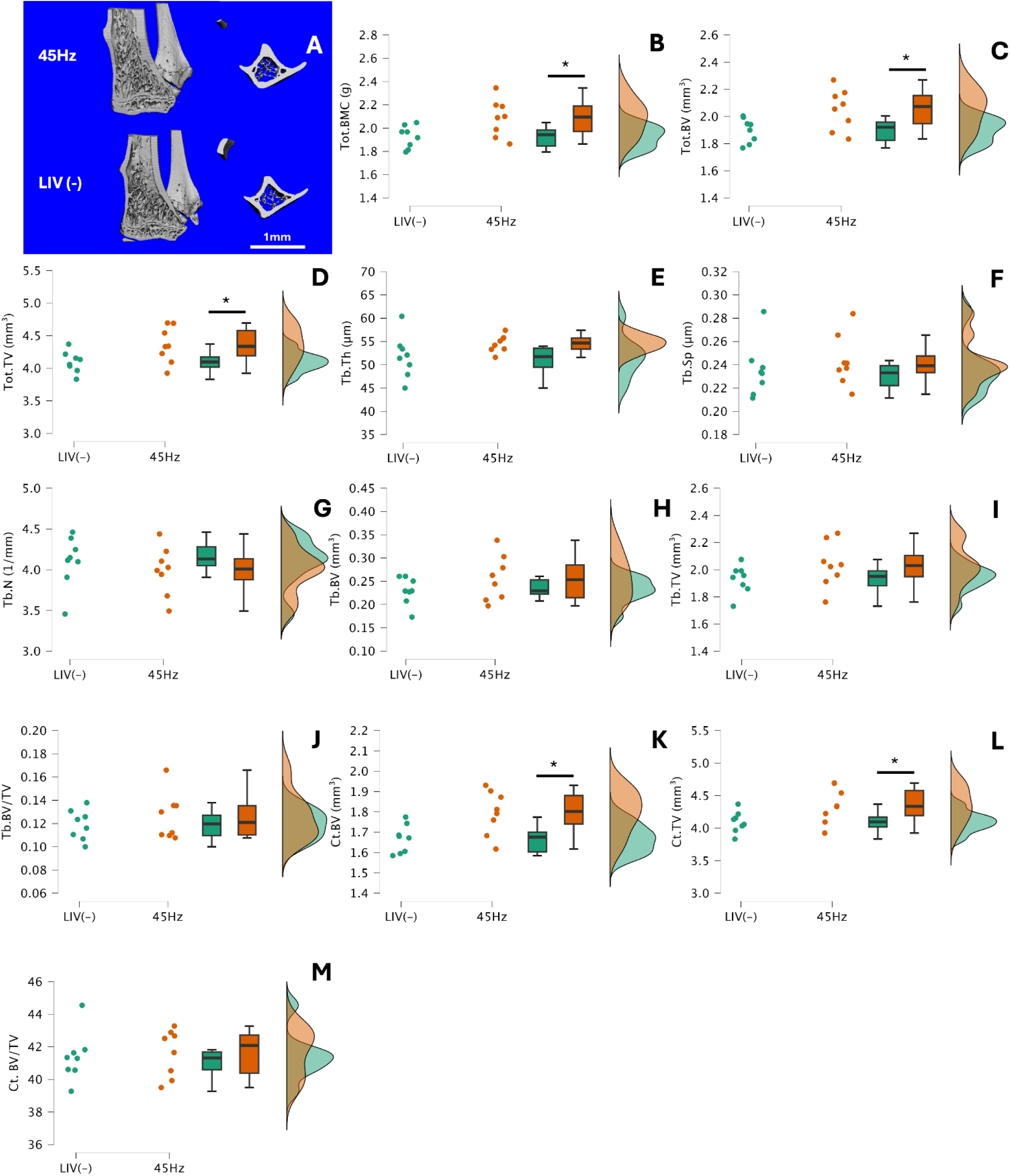
μCT measures derived from tibias of 24weeks old C57BL/6 mice subjected to LIV regime. (A) Representative 3D reconstruction of proximal metaphyseal trabecular region of interest (coronal cut) and mid-shaft cortical cross-section of the tibia’s region of interest; (B) Total Bone Mineral Content; (C) Total Bone Volume; (D) Total Tissue Volume, (E) Trabecular Thickness; (F) Trabecular Spacing; (G) Trabecular Number; (H) Trabecular Bone Volume; (I) Trabecular Tissue Volume ; (J) Trabecular Bone Fraction; (K) Cortical Bone Volume; (L) Cortical Tissue Volume; (M) Cortical Bone Fraction; (N) Cortical Area; (O) Cortical Thickness. Significance between groups tested using T-Student test or Mann-Whitney-U test when not parametric, *p< 0.05, ** p< 0.01

## DISCUSSION

In this study, we systematically reviewed the existing evidence on LIV protocols for the improvement of BV/TV in small rodents, ranked the efficacy of distinct frequency-intensity combination by using a Bayesian network meta-analysis, and performed an experimental confirmation for a distinguished frequency-intensity combination. Besides the individual limitations in each of the included studies, our primary findings reveal that LIV produces non-uniform responses among hindlimb long bones, as the tibia and femur exhibited different responses to similar LIV parameters. The LIV protocol of 45Hz at 2g exhibited the highest and significant probability of increasing BV/TV for the tibia, while for the femur, the frequency-intensity combination that ranked first on chances of effective results was 90Hz at 1.4g. Even so, effects on femur were not as pronounced as those found with tibia, as none of the interventions done on femur showed a significant effect. These site-specific results suggest that LIV may be tuned by factors like bone tissue properties, bone morphology, and habitual loading patterns. As LIV generates small acellerations on the bone tissue, on the fluids within the bone like bone marrow^7–9^ and on the interstitial fluid within the lacuno-canalicular system^56,57^, the fluid shear ellicts mechanotransductions on osteocytes^10,56^ which in turn evoke a complex downstream signaling for osteogenic responses^58^. Therefore, factors like the imposed acceleration, fluid shear and the local bone properties may contribute to the osteogenic responses in response to LIV.

As for the different frequency and acceleration protocols analysed, our Bayesian network meta-analysis results suggest that for the tibia, as a primary weight-loaded bone, a mid-range frequency (45Hz at 2g) may cause a significant BV/TV increase. LIV at 45Hz interventions not only performed substantially different intervention using other frequencies, but also ranked with the highest chances to provide the best BV/TV response. Regarding acceleration, while most reported LIV protocols used accelerations up to 2g, rodent running can elicit up to 1g of acceleration on the same region^59^. According to *in silico* results from fluid shear stress at bone - marrow interface, accelerations between 0.1 and 2g can create shear stresses up to 2Pa^9,60,61^, which is enough to mechanically stimulate osteoblasts, osteoclasts and mesenchymal cells within the bone^60^.

Given that strain magnitude or rate by itself are the main factors impacting the velocity and shear stress around osteocytes located inside canaliculi^56,57^, the observed higher responses in the tibia when using 2g instead of 0.2g for the same frequency (45Hz), may be justified by the increased strain or fluid shear signaling. Our μCT results even confirms that under 0.2g, LIV set to 45Hz was enough to elicity significant increases on tibia bone mineral content, on cortical bone volume, and on cortical tissue volume.

In terms of femoral responses, our Baeysian network meta-analysis showed the intervention with highest possibility to perform better presents a frequency twice as large (90Hz), with roughly 75% of the acceleration (1.4g) as the one that performed best for tibia. On the rankogram, the LIV6 setup at 90Hz and 1.4g was followed by the same two LIV setups that performed significantly better for tibia, 45Hz at 2g and 45Hz at 0.2g. Despite these three setups being highlighted, none of the investigated LIV setups in this paper presented significant improvement for BV/TV for femurs. On the other hand, our μCT results reinforce that 45Hz at 0.2g benefits femur microstructure, by presenting significant increases in many of the observed variables. It is important to emphasize that we analysed studies including both rats and mice. According to the literature, rats tend to present a less variable bone turnover compared to mice^62^. In our study, four out of the five included studies analyzing femoral BV/TV were conducted in rats. Although Bayesian network meta-analysis can account for between-study heterogeneity, we cannot ignore that the composition of the included studies may have influenced the observed findings regarding femoral responses to LIV. Combined, this may be the reason we observed significant increases in the femur μCT analysis but not on the femurs in the networks meta-analsysis.

Although, despite the variety of combinations of frequency, acceleration and animal model observed in our study, different responses between femur and tibia may be expected even when using a stardardized vibration, acceleration and animal model. We believe that a divergence can occur due to combination of variables such as site-specific physiology, mechanisms like attenuated accelerations, and knee biomechanics.

Previous reports have show that the femur seems to have a not only a lower turnover rate^63^, but also lower trabecular bone proportion compared to tibia^37,64^. Additionally, previous findings led us to believe that femur responses to LIV might be more efficient at higher frequencies. Despite the rodent anatomy and dynamic stance potentially attenuating mechanical signals between adjacent bone segments through soft tissue damping, muscle activity damping and dynamic joint angles damping^65^, the review by Reynols et al. suggests that resonance might occur in whole body vibration at frequencies between 80 to 90 Hz when applied to mice^66^. Given the anatomical similarities in small rodent model, the LIV imposed at 90Hz and 1.4g might scored better for femur given that frequencies below 90Hz might not be sufficient to transmit sufficient energy beyond the knee joint.

Despite pointing out which LIV setup is likely to perform better in small rodents, our findings also indicate that using a LIV protocol as “one-size-fits-all” might not be the best practice. Also, besides the results suggesting that the response to LIV may be site/bone dependent, our results only account for experiments with mice and rats with no additional health conditions. Therefore, it is reasonable to assume that, depending on the experiment model (bone fracture, osteoporosis, disuse-induced bone loss), interventions might require a specific frequency and/or intensity optimization. LIV has been widely tested and used in humans, and while the underlying principle-specific optimization is likely to be preserved, translating our findings to human application is unlikely to produce similar results. Although results from rodent model studies are highly translational for human application^67,68^, the normal stance mechanics are widely differing between species, affecting muscle activity^69^, and damper effect^70^ amongst others not mentioned here. We suggest that future experimental studies should consider including multiple frequency-intensities when aiming to identify the most effective protocols for human skeletal sites that are susceptible to bone conditions such as osteoporosis, osteopenia and bone fractures. The use of Bayesian inference provides a rigorous statistical foundation for our analysis. Employing it in our network meta-analysis, enabled us to combine published data and provide a probabilistic ranking of all included LIV regimens, offering a broader guide for parameter selection when compared to traditional pairwise methods. Combined with surface under the cumulative ranking, it further contributes to drawing conclusions from quantitative measures for such ranking. We further reinforce our findings by analyzing potential bias within the studies. Still, our study presents limitations when it comes to included studies and residual confounding. Given our eligibility criteria, the mild number of seven selected studies limits the external validity of our findings. Also, the fact that network meta-analysis accommodates for heterogeneities like rodent strains, age, LIV protocol and outcome report, does not mean that the residual confounding should be fully neglected.

In conclusion, our systematic review and Bayesian meta-analysis, along with the experimental studies, demonstrate that the osteogenic response to LIV is dependent on both frequency-intensity and anatomical site (bone dependent). Besides being closer to the vibration source, the tibia may resonate better morphological responses to lower frequencies (45Hz) compared to the femur’s preferred frequency (90Hz). The finding presented in this study highlight the importance of vibration parameter optimization and bone target specificity while designing mechanical stimulation protocols for eithertbone adaptation or prevention of ongoing challenges in bone disorders.

## FUNDING STATEMENT

This study was supported by AG059923, P20GM109095, NSF 2025505, and NSF 2431083.

## ABREVIATIONS LIST

LIV: Low intensity vibration
g: Gravitational acceleration
Hz: Hertz
CrI: Credible Interval
ms: millisecond
Pa: Paschal
BV/TV: Bone volume fraction
µCT: Micro computed tomography
µε: Micro strain
NMA: Metwork Meta-Analysis
MCMC: Markov chain Monte Carlo

## Notes

### Competing Interest Statement

The authors have declared no competing interest.

