## Supplemental table and figures for "Vibration’s frequency and intensity for optimal setup for enhancement bone response in small rodents: A systematic review and Bayesian network meta-analysis"

### SUPPLEMENTAL MATERIAL

### *DATABASES SEARCH STRATEGIES (Key words and Mesh terms)*

**PubMed Search Strategy:** [mh] mesh terms; [tw] text words; [tiab] title and abstract; [all] all fields; * Replaces multiple characters at the end of a root word

| Concept 1<br>Population | Concept 2<br>Intervention | Concept 3<br>Comparison | Concept 4<br>Outcome |
|---|---|---|---|
| Mouse [tw] (#1) | Low intensity vibration [tw] (#41) | BV/TV [tw] (#61) | Bone mass [all] (#83) |
| Mice [mh] (#2) | Whole body vibration [tw] (#42) | B.V./T.V. [all] (#62) | Bone quality [all] (#84) |
| Mus musculus domesticus [tw] (#3) | Vibration training [tw] (#43) | BV./TV. [tw] (#63) | Bone volume [all] (#85) |
| C57BL/6J [tw] (#4) | Low frequency vibration [tw] (#44) | Bone volume tissue volume [tw] (#64) | Bone density [all] (#86) |
| C57BL/6 [tw] (#5) | High frequency vibration [tw] (#45) | BV.f. [all] (#65) | |
| C57BL [tw] (#6) | LIV [tw] (#46) | BVf [all] (#66) | |
| C57 black 6 [tw] (#7) | Vibration platform [tw] (#47) | Bone volume fraction [tw] (#67) | |
| B6 [tw] (#8) | Vibration intervention [tw] (#48) | Tibia* [tw] (#68) | |
| C57 [tw] (#9) | Low magnitude high frequency vibration [tw] (#49) | Femur [tw] (#69) | |
| Black 6 [tw] (#10) | LMHFV [tw] (#50) | Femor*[tw] (#70) | |
| BALB/cByJ [tw] (#11) | Low magnitude mechanical signals [tw] (#51) | Micro-CT [tw] (#71) | |
| BALB/c [tw] (#12) | LMMS [tw] (#52) | Micro-computed tomography [tw] (#72) | |
| BALB C mice [tw] (#13) | Low-intensity whole-body vibration [tw] (#53) | X-Ray Microtomography [all] (#73) | |
| DBA/2J [tw] (#14) | LIWBV [tw] (#54) | Microcomputed tomography [tw] (#74) | |
| Mice, Inbred CBA [mh] (#15) | low-magnitude high-frequency [tw] (#55) | Micro computed tomography [tw] (#75) | |
| Swiss Webster [tw] (#16) | LMHF [tw] (#56) | CT scan* [tw] (#76) | |
| Mice, Inbred Strains [mh] (#17) | WBV [tw] (#57) | X-ray CT [tw] (#77) | |
| Rats [mh] (#18) | Low-amplitude whole-body vibration [tw] (#58) | | |
| Rat [tw] (#19) | low-magnitude high frequency mechanical signals [all] (#59) | | |
| Rats, Inbred Strains [mh] (#20) | | | |

| | | | |
|---|---|---|---|
| Rattus norvegicus [tw] (#21) | | | |
| Rats, Sprague-Dawley [mh] (#22) | | | |
| Rats, Inbred F344 [mh] (#23) | | | |
| black hooded Lister rats [tw] (#24) | | | |
| hooded Lister rats [tw] (#25) | | | |
| black-hooded Lister rats [all] (#26) | | | |
| Rats, Long-Evans [mh] (#27) | | | |
| Norway rat [tw] (#38) | | | |
| Rats, Wistar [mh] (#29) | | | |
| **#30 =** (#1 OR #2 OR #3 OR #4 OR #5 OR #6 OR #7 OR #8 OR #9 OR #10 OR #11 OR #12 OR #13 OR #14 OR #15 OR #16 OR #17 OR #18 OR #19 OR #20 OR #21 OR #22 OR #23 OR #24 OR #25 OR #26 OR #27 OR #28 OR #29 | **#50 =** (#31 OR #32 OR #33 OR #34 OR #35 OR #36 #37 OR #38 OR #39 OR #40 OR #41 OR #42 OR #43 OR #44 OR #45 OR #46 OR #47 OR #48 OR #49) | **#68** = (#51 OR #52 OR #53 OR #54 OR #55 OR #56 OR #57 OR #58 OR #59 OR #60 OR #61 OR #62 OR #63 OR #64 OR #65 OR #66 OR #67 | **#73** = (#69 OR #70 OR #71 OR #72) |
| **#74 = #30 AND #50 AND #68 AND #73 = 84 results as it 18/04/2025** | | | |

**RETURNS:** 84;

**TOTAL RETURNS, separated columns:** POPULATION 3,730,062; INTERVENTION 5,817; COMPARISON 513,674; OUTCOME 107,476;

**TOTAL SUM OF RETURNS:** 4,357,029.

**Web of Science search strategy:** [all] = all text words; [ti] = title; * Replaces multiple characters at the end of a root word

| Concept 1<br><br>Population | Concept 2<br><br>Intervention | Concept 3<br><br>Comparison | Concept 4<br><br>Outcome |
|---|---|---|---|
| Mouse [all] (#1) | Low intensity vibration [all] (#34) | BV/TV [all] (#54) | Bone mass [all] (#72) |
| Mice [all] (#2) | Whole body vibration* [all] (#35) | B.V./T.V. [all] (#55) | Bone quality [all] (#73) |
| Mus musculus domesticus [all] (#3) | Vibration training [all] (#36) | BV./TV. [all] (#56) | Bone volume [all] (#74) |
| C57BL/6J [all] (#4) | Low frequency vibration [all] (#37) | Bone volume tissue volume [all] (#57) | Bone density [all] (#75) |
| C57BL/6 [all] (#5) | High frequency vibration [all] (#38) | BV.f. [all] (#58) | |
| C57BL [all] (#6) | LIV [all] (#39) | BVf [all] (#59) | |
| C57 black 6 [all] (#7) | Vibration platform [all] (#40) | Bone volume fraction [all] (#60) | |
| B6 [all] (#8) | Vibration intervention [all] (#41) | Tibia* [all] (#61) | |
| C57 [all] (#9) | Low magnitude high frequency vibration [all] (#42) | Femur [all] (#62) | |
| Black 6 [all] (#10) | LMHFV [all] (#43) | Femor* [all] (#63) | |
| BALB/cByJ [all] (#11) | Low magnitude mechanical signals [all] (#44) | Micro-CT [all] (#64) | |
| BALB/c [all] (#12) | LMMS [all] (#45) | Micro-computed tomography [all] (#65) | |
| BALB C mice [all] (#13) | Low-intensity whole-body vibration [all] (#46) | X-Ray Microtomography [all] (#66) | |
| BALB/cByJ (BALB) | LIWBV [all] (#47) | Microcomputed tomography [all] (#67) | |
| DBA/2J [all] (#15) | Low-magnitude high-frequency [all] (#48) | Micro computed tomography [all] (#68) | |
| CBA [all] (#16) | LMHF [all] (#49) | CT scan* [all] (#69) | |
| Swiss Webster [all] (#17) | WBV [all] (#50) | X-ray CT [all] (#70) | |
| Mice, Inbred Strains [all] (#18) | Low-amplitude whole-body vibration [all] (#51) | | |
| Rats [all] (#19) | Low-magnitude high frequency mechanical signals [all] (#52) | | |
| Rat [all] (#20) | | | |
| Rats, Inbred Strains [all] (#21) | | | |
| Rattus norvegicus [all] (#22) | | | |

| | | | |
|---|---|---|---|
| Sprague-Dawley [all] (#23) | | | |
| F344 [all] (#24) | | | |
| Lister black hooded [all] (#25) | | | |
| black hooded Lister rats [all] (#26) | | | |
| hooded Lister rats [all] (#27) | | | |
| black-hooded Lister rats [all] (#28) | | | |
| Long-Evans [all] (#30) | | | |
| Norway [all] (#31) | | | |
| Wistar rats [all] (#32) | | | |
| **#33=** #1 OR #2 OR #3 OR #4 OR #5 OR #6 OR #7 OR #8 OR #9 OR #10 OR #11 OR #12 OR #13 OR #14 OR #15 OR #16 OR #17 OR #18 OR #19 OR #20 OR #21 OR #22 OR #23 OR #24 OR #25 OR #26 OR #27 OR #28 OR #29 OR #30 OR #31 OR #32 | **#53 =** #34 #35 #36 #37 OR #38 OR #39 OR #40 OR #41 OR #42 OR #43 OR #44 OR #45 OR #46 OR #47 OR #48 OR #49 OR #50 OR #51 OR #52 | **#71** = #54 #55 #56 #57 OR #58 OR #59 OR #60 OR #61 OR #62 OR #63 OR #64 OR #65 OR #66 OR #67 OR #68 OR #69 OR #70 | **#76** = (#72 OR #73 OR #74 OR #75) |
| **#77 = #33 AND #53 AND #71 AND #76 = 80 results as it 18/APR/2025** | | | |

**RETURNS: 80;**

**TOTAL RETURNS, separated columns:** POPULATION 4,513,919; INTERVENTION 22,286; COMPARIZON 454,133; OUTCOME 63,056;

**TOTAL SUM OF RETURNS:**4,643,394.

**EMBASE search strategy:** xxx= all fields; [:ti]= title; * Replaces multiple characters at the end of a root word

| Concept 1<br><br>Population | Concept 2<br><br>Intervention | Concept 3<br><br>Comparison | Concept 4<br><br>Outcome |
|---|---|---|---|
| Mouse (#1) | Low intensity vibration* (#34) | BV/TV (#54) | Bone mass (#72) |
| Mice (#2) | Whole body vibration (#35) | B.V./T.V. (#55) | Bone quality (#73) |
| Mus musculus domesticus (#3) | Vibration training (#36) | BV./TV. (#56) | Bone volume (#74) |
| C57BL/6J (#4) | Low frequency vibration (#37) | Bone volume tissue volume (#57) | Bone density (#75) |
| Inbred C57BL (#5) | High frequency vibration (#38) | BV.f. (#58) | |
| C57 black 6 (#6) | LIV (#39) | BVf (#59) | |
| B6 (#7) | Vibration intervention (#40) | Bone volume fraction (#60) | |
| C57 (#8) | Vibration platform (#41) | Tibia* (#61) | |
| Black 6 (#9) | Low magnitude high frequency vibration (#42) | Femur (#62) | |
| BALB/cByJ (#10) | LMHFV (#43) | Femor* (#63) | |
| BALB/c (#11) | Low magnitude mechanical signals (#44) | Micro-CT (#64) | |
| BALB C mice (#12) | LMMS (#45) | Micro-computed tomography (#65) | |
| BALB/cByJ (BALB) (#13) | Low-intensity whole-body vibration (#46) | X-Ray Microtomography (#66) | |
| DBA/2J (#14) | LMHF (#47) | Microcomputed tomography (#67) | |
| CBA mice (#15) | WBV (#48) | Micro computed tomography (#68) | |
| Swiss Webster mouse (#16) | Low-amplitude whole-body vibration (#49) | CT scan* (#69) | |
| Inbred mouse Strain (#17) | low-magnitude high frequency mechanical signals (#50) | X-ray CT (#70) | |
| Rat (#18) | LIWBV (#51) | | |
| Rats (#19) | low-magnitude high-frequency (#52) | | |
| Inbred rat strain (#20) | | | |
| Rattus norvegicus (#21) | | | |
| Sprague-Dawley rat (#22) | | | |
| F344 (#23) | | | |
| Lister black hooded (#24) | | | |
| C57BL/6 (#25) | | | |
| black hooded Lister rats (#26) | | | |
| hooded Lister rats (#27) | | | |
| black-hooded Lister rats (#28) | | | |

| | | | |
|---|---|---|---|
| Long Evans rat (#30) | | | |
| Norway rat (#31) | | | |
| Wistar rat (#32) | | | |
| **#33 =** (#1 OR #2 OR #3 OR #4 OR #5 OR #6 OR #7 OR #8 OR #9 OR #10 OR #11 OR #12 OR #13 OR #14 OR #15 OR #16 OR #17 OR #18 OR #19 OR #20 OR #21 OR #22 OR #23 OR #24 OR #25 OR #26 OR #27 OR #28 OR #29 OR #30 OR #31 OR #32) | **#53 =** (#34 OR #35 #37 OR #38 OR #39 OR #40 OR #41 OR #42 OR #43 OR #44 OR #45 OR #46 OR #47 OR #48 OR #49 OR #50 OR #51 OR #52) | **#71** = (#54 OR #55 OR #56 OR #57 OR #58 OR #59 OR #60 OR #61 OR #62 OR #63 OR #64 OR #65 OR #66 OR #67 OR #68 OR #69 OR #70) | **#76** = (#72 OR #73 OR #74 OR #75) |
| **#77 = #33 AND #53 AND #71 AND #76 = 148 results as it 18/APR/2025** | | | |

**RETURNS:** 148;

**TOTAL RETURNS, separated columns:** POPULATION 5,106,728; INTERVENTION 24,878; COMPARISON 839,343; OUTCOME 172,853;

**TOTAL SUM OF RETURNS:** 5,393,802.

**CINAHL (Ebsco) search strategy:** xxx = all fields; [TI] = title; * Replaces multiple characters at the end of a root word

| **Concept 1**<br><br>**Population** | **Concept 2**<br><br>**Intervention** | **Concept 3**<br><br>**Comparison** | **Concept 4**<br><br>**Outcome** |
|---|---|---|---|
| Mouse | Low intensity vibration | Tibia* | Bone mass |
| Mice | Whole body vibration | Femur | Bone quality |
| Mus musculus domesticus | Vibration training | Femor* | Bone volume |
| C57BL/6J | Low frequency vibration | Micro-CT | Bone density |
| C57BL/6 | High frequency vibration | Micro-computed tomography | |
| C57BL | LIV | X-Ray Microtomography | |
| C57 black 6 | Vibration platform | Microcomputed tomography | |
| B6 | Vibration intervention | BV/TV | |
| C57 | Low magnitude high frequency vibration | B.V./T.V. | |
| Black 6 | LMHFV | BV./TV. | |
| BALB/cByJ | Low magnitude mechanical signals | Bone volume tissue volume | |
| BALB/c | LMMS | BV.f. | |
| BALB C mice | Low-intensity whole-body vibration | BVf | |
| BALB/cByJ (BALB) | LIWBV | Bone volume fraction | |
| DBA/2J | low-magnitude high-frequency | Micro computed tomography | |
| CBA | LMHF | CT scan* | |
| Swiss Webster | WBV | X-ray CT | |
| Inbred Mice Strain | Low-amplitude whole-body vibration | | |
| Rats | Low-magnitude high frequency mechanical signals | | |
| Rat | | | |
| Inbred Strain rat | | | |
| Rattus norvegicus | | | |
| Sprague-Dawley | | | |
| F344 | | | |
| Lister black hooded | | | |
| black hooded Lister rats | | | |
| hooded Lister rats | | | |
| black-hooded Lister rats | | | |
| Long Evans rat | | | |
| Norway rats | | | |
| Wistar rats | | | |
| Mice | | | |

<table>
<tr>
<td>(mouse OR mice OR mus musculus domesticus OR c57bl/6j OR c57bl/6 OR c57bl OR c57 black 6 OR b6 OR c57 OR balb/cbyj OR balb/c OR balb c mice OR balb/cbyj (balb) OR dba/2j OR cba OR swiss webster OR inbred strain mice OR rats OR rat OR inbred strain rat OR rattus norvegicus OR sprague-dawley OR f344 OR lister black hooded OR black hooded lister rats OR hooded lister rats OR black-hooded lister rats OR long-evans OR norway OR wistar rats NOT TI rabbit NOT TI rabbits NOT TI turkey NOT TI cricetidae NOT TI guinea pigs NOT TI guinea pig NOT TI horses NOT TI horse NOT TI dogs NOT TI dog)</td>
<td>low intensity vibration OR whole body vibration* OR vibration training OR low frequency vibration OR high frequency vibration OR liv OR vibration platform OR vibration intervention OR low magnitude high frequency vibration OR lmhfv OR low magnitude mechanical signals OR lmms OR low-intensity whole-body vibration OR liwbv OR low-magnitude high-frequency OR lmhf OR wbv OR low-amplitude whole-body vibration OR low-magnitude high frequency mechanical signals)</td>
<td>(bv/tv OR b.v./t.v. OR bv./tv. OR bone volume tissue volume OR bv.f. OR bvf OR bone volume fraction OR tibia* OR femur OR femor* OR micro-ct OR micro-computed tomography OR x-ray microtomography OR microcomputed tomography OR micro computed tomography OR ct scan* OR x-ray ct NOT osteotomy NOT fracture NOT fracture site NOT callus)</td>
<td>(bone mass OR bone quality OR bone volume OR bone density)</td>
</tr>
<tr>
<td colspan="4">(mouse OR mice OR mus musculus domesticus OR c57bl/6j OR c57bl/6 OR c57bl OR c57 black 6 OR b6 OR c57 OR balb/cbyj OR balb/c OR balb c mice OR balb/cbyj (balb) OR dba/2j OR cba OR swiss webster OR inbred strain mice OR rats OR rat OR inbred strain rat OR rattus norvegicus OR sprague-dawley OR f344 OR lister black hooded OR black hooded lister rats OR hooded lister rats OR black-hooded lister rats OR long-evans OR norway OR wistar rats NOT TI rabbit NOT TI rabbits NOT TI turkey NOT TI cricetidae NOT TI guinea pigs NOT TI guinea pig NOT TI horses NOT TI horse NOT TI dogs NOT TI dog) AND (low intensity vibration OR whole body vibration* OR vibration training OR low frequency vibration OR high frequency vibration OR liv OR vibration platform OR vibration intervention OR low magnitude high frequency vibration OR lmhfv OR low magnitude mechanical signals OR lmms OR low-intensity whole-body vibration OR liwbv OR low-magnitude high-frequency OR lmhf OR wbv OR low-amplitude whole-body vibration OR low-magnitude high frequency mechanical signals) AND (bv/tv OR b.v./t.v. OR bv./tv. OR bone volume tissue volume OR bv.f. OR bvf OR bone volume fraction OR tibia* OR femur OR femor* OR micro-ct OR micro-computed tomography OR x-ray microtomography OR microcomputed tomography OR micro computed tomography OR ct scan* OR x-ray ct NOT osteotomy NOT fracture NOT fracture site NOT callus) AND (bone mass OR bone quality OR bone volume OR bone density)<br>= 25 results as it 18/APR/2025</td>
</tr>
</table>

**RETURNS:** 25;

**TOTAL RETURNS, separated columns:** POPULATION 221,153; INTERVENTION 2,418; COMPARIZON 113,314; OUTCOME 33,517;

**TOTAL SUM OF RETURNS:** 370,402.

### *ELIGIBILITY CRITERIA*

During full reading of the selected papers, we used the following hieraquic excluding criteria to exclude studies that did not match the variables we aimed to analyse: $1^{st}$ Study did not use mice or rats; $2^{nd}$ Study used non mature animals (12 weeks for mice <21 weeks for rats); $3^{rd}$ Study involving elderly animals (≥ 73 weeks for mice ≥ 101 weeks for rats); $4^{th}$ Study did not vibrate the whole animal; $5^{th}$ Study did not investigate bones; $6^{th}$ Study did not did not look at limbs' bones; $7^{th}$ Study did not look at bone microarchitecture (BV/TV or BV.f); $8^{th}$ Study evaluated callus areas (intentional fractures or implants); $9^{th}$ Study used LIV combined with other variables; $10^{th}$ Study was not a peer reviewed or was an abstract; $11^{th}$ Other reasons (Sample too small; Micro-CT nomenclature was not standard; Study does not present control group for LIV; LIV intensity or frequency was not disclosed; Study used frequencies <10Hz or >100Hz).

**Supplementary Figure 1A:** Fixed effects model vs. random effects model for tibia dataset

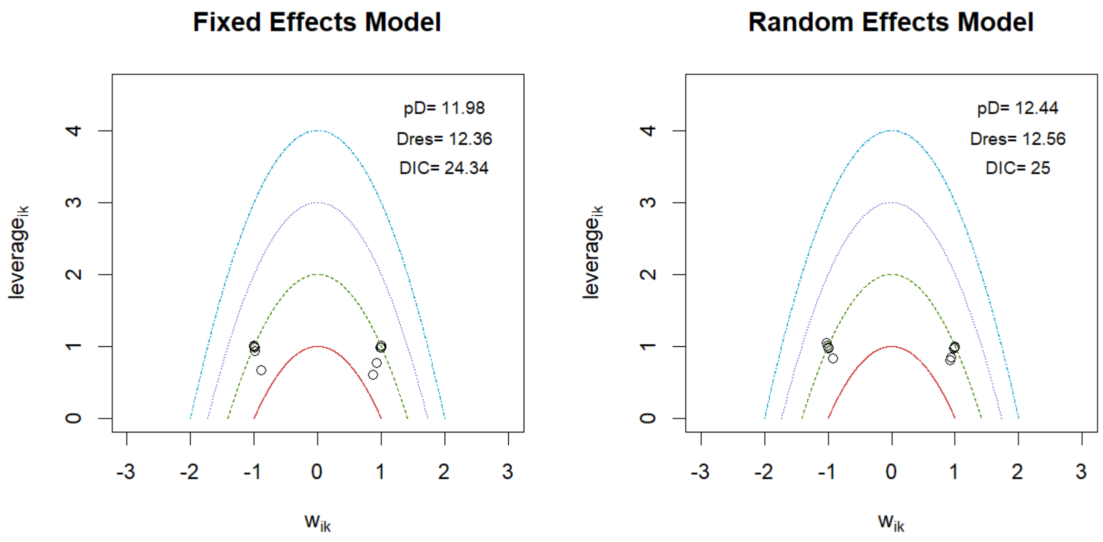


**Supplementary Figure 1B:** Fixed effects model vs. random effects model for femur dataset

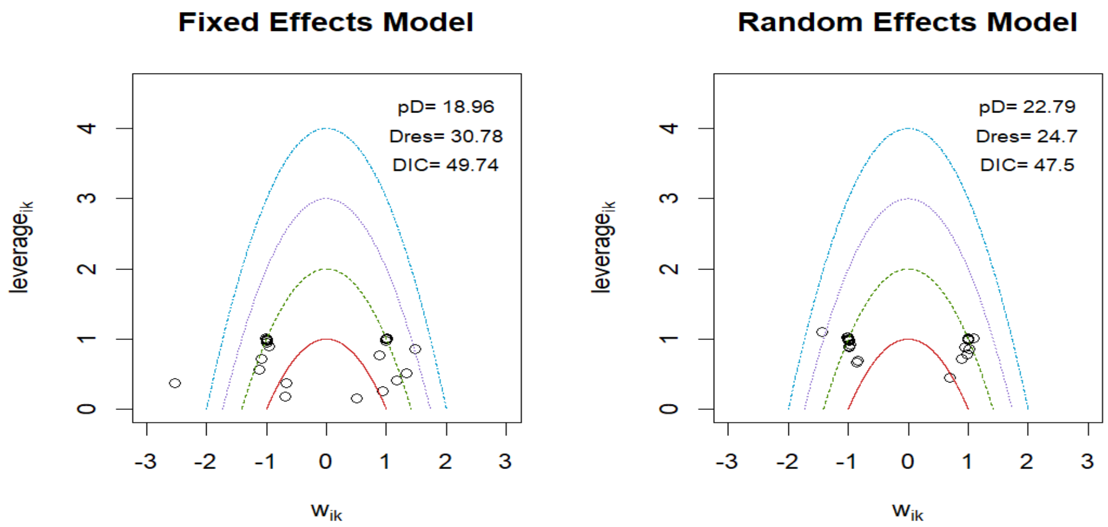

**Supplementary Figure 2A:** Trace and Density Plots for tibia dataset

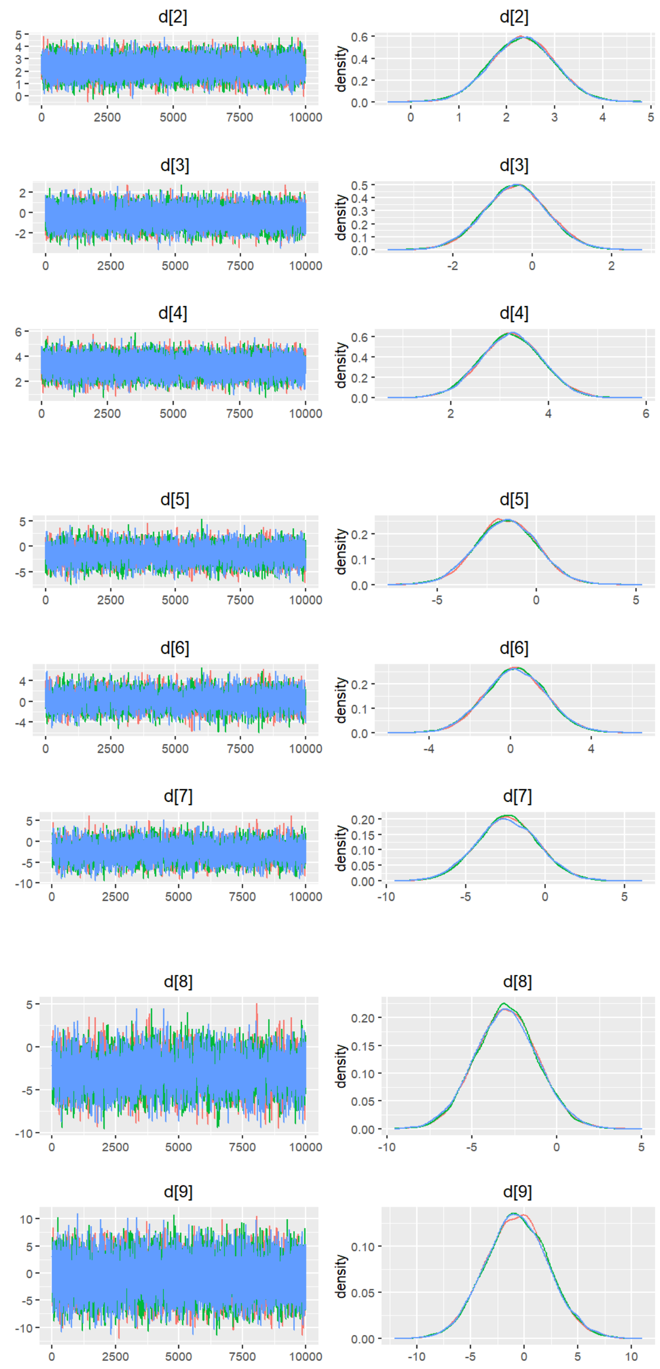

**Supplementary Figure 2B:** Trace and Density Plots for femur dataset

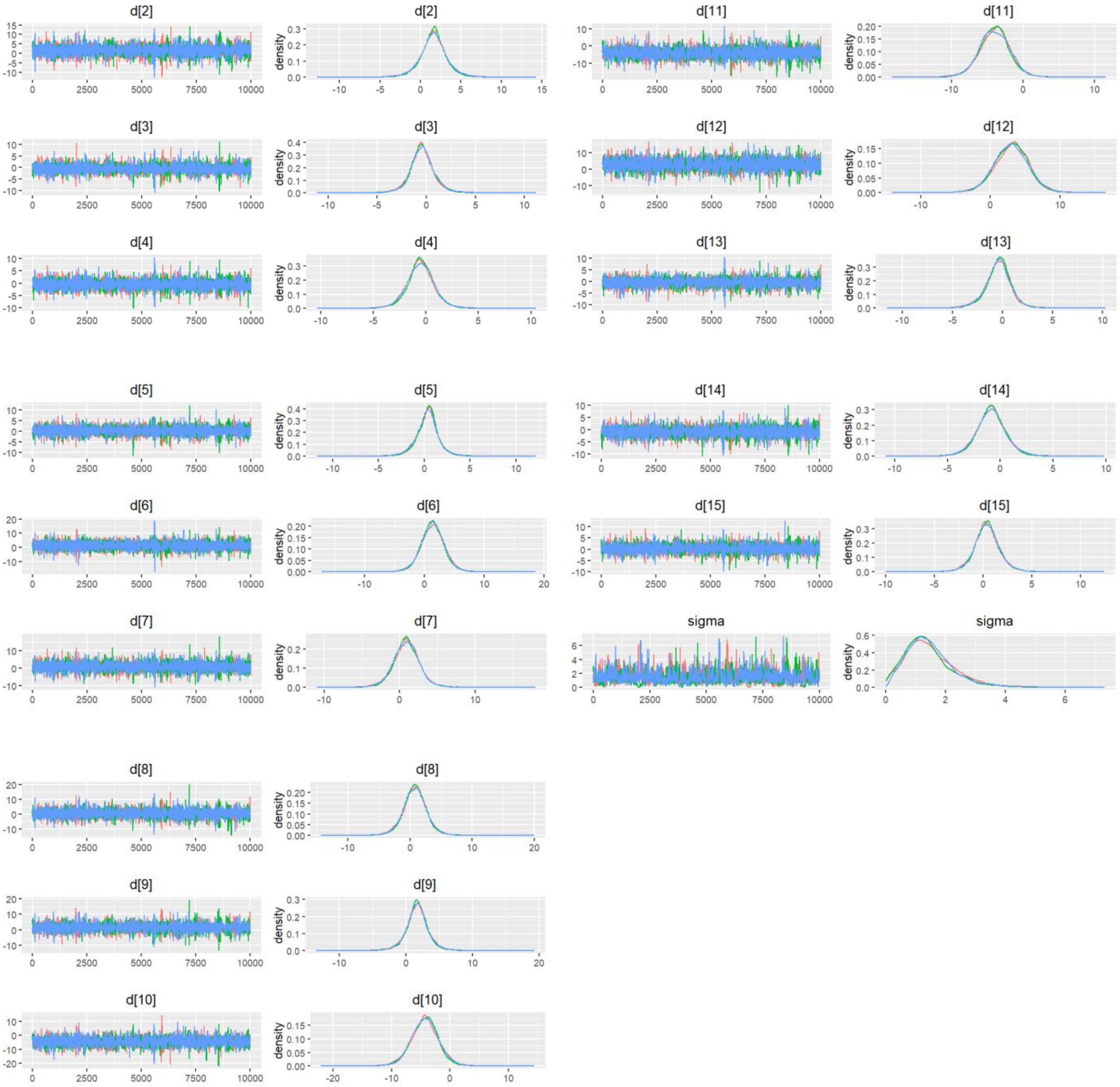

**Supplementary Figure 3A:** Inconsistency Plot for Tibia analysis

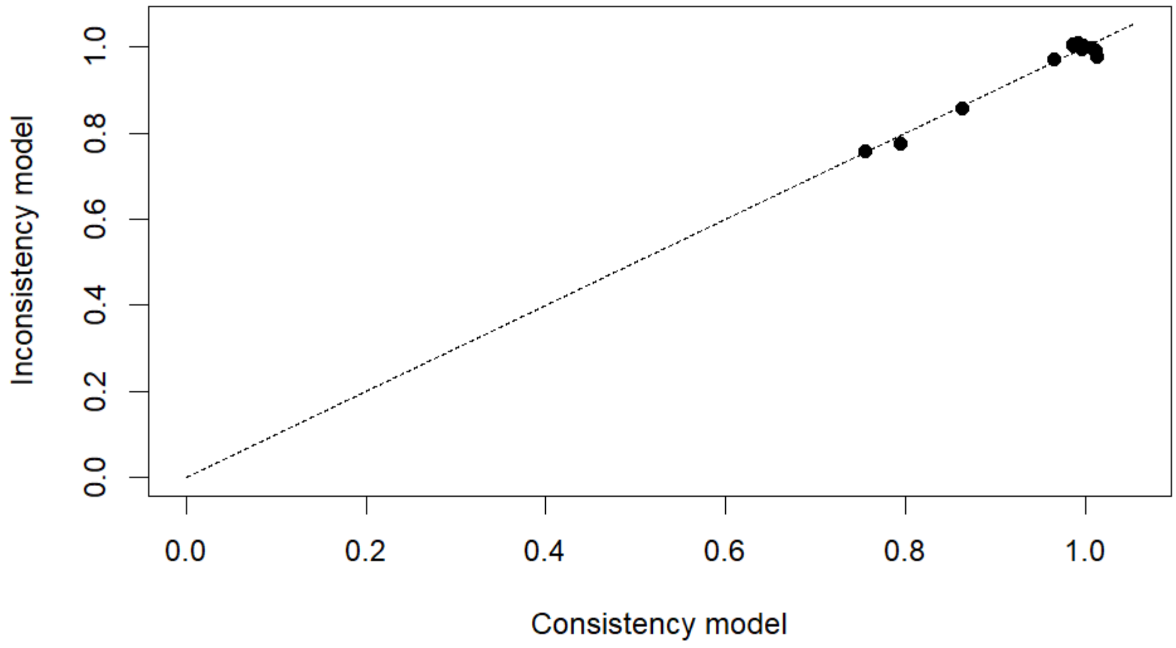


**Supplementary Figure 3B:** Inconsistency Plot for Femur analysis

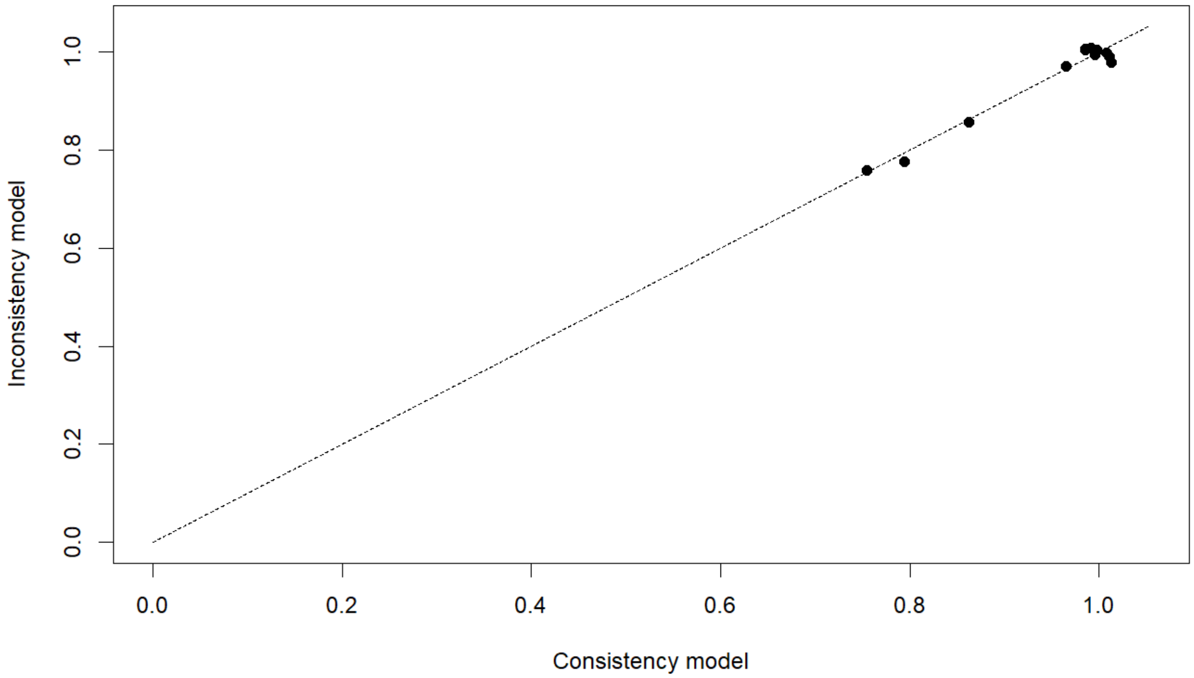

**Supplementary Figure 4A:** SUCRA plot of Tibia LIV levels

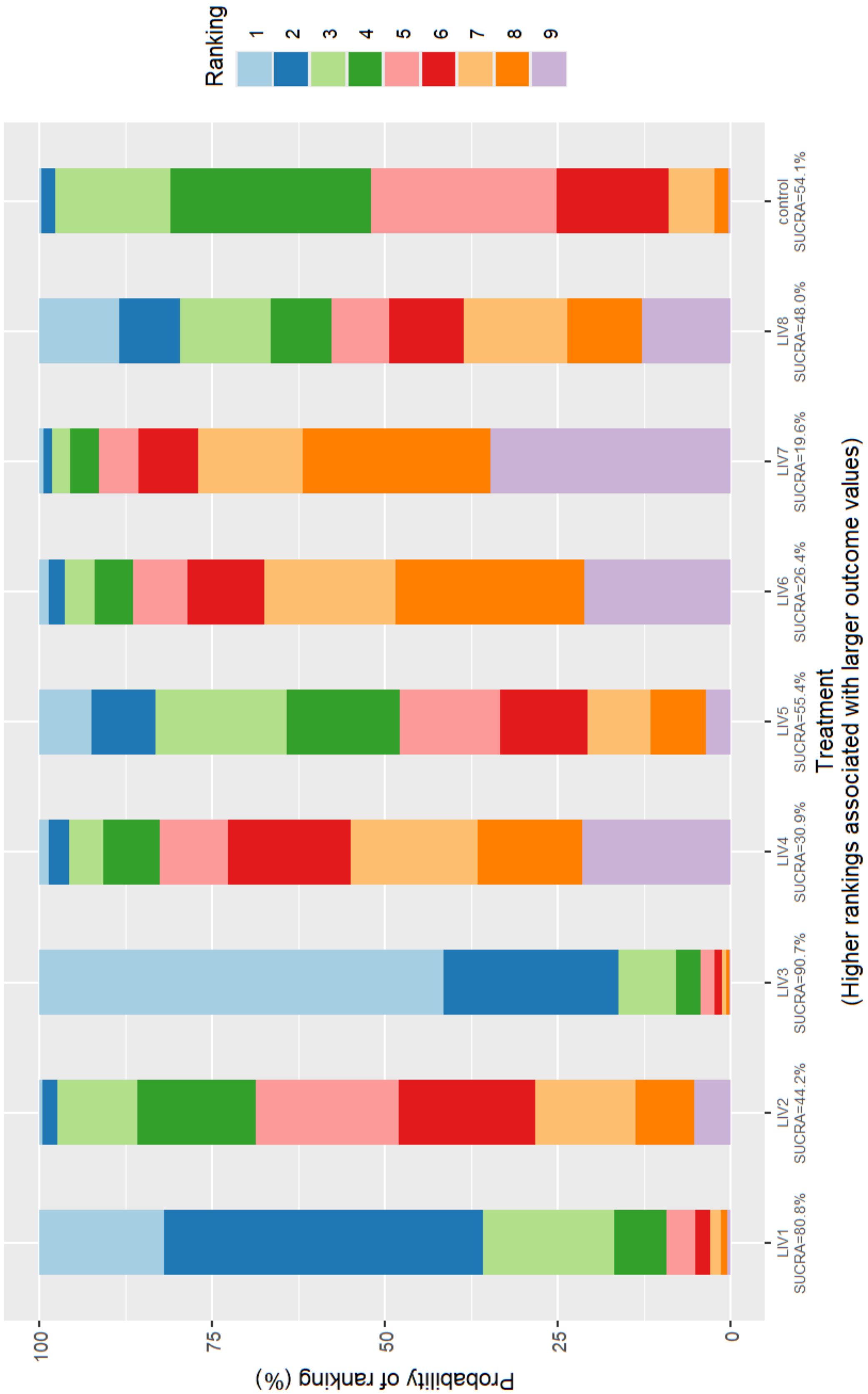

**Supplementary Figure 4B:** SUCRA plot of Femur LIV levels

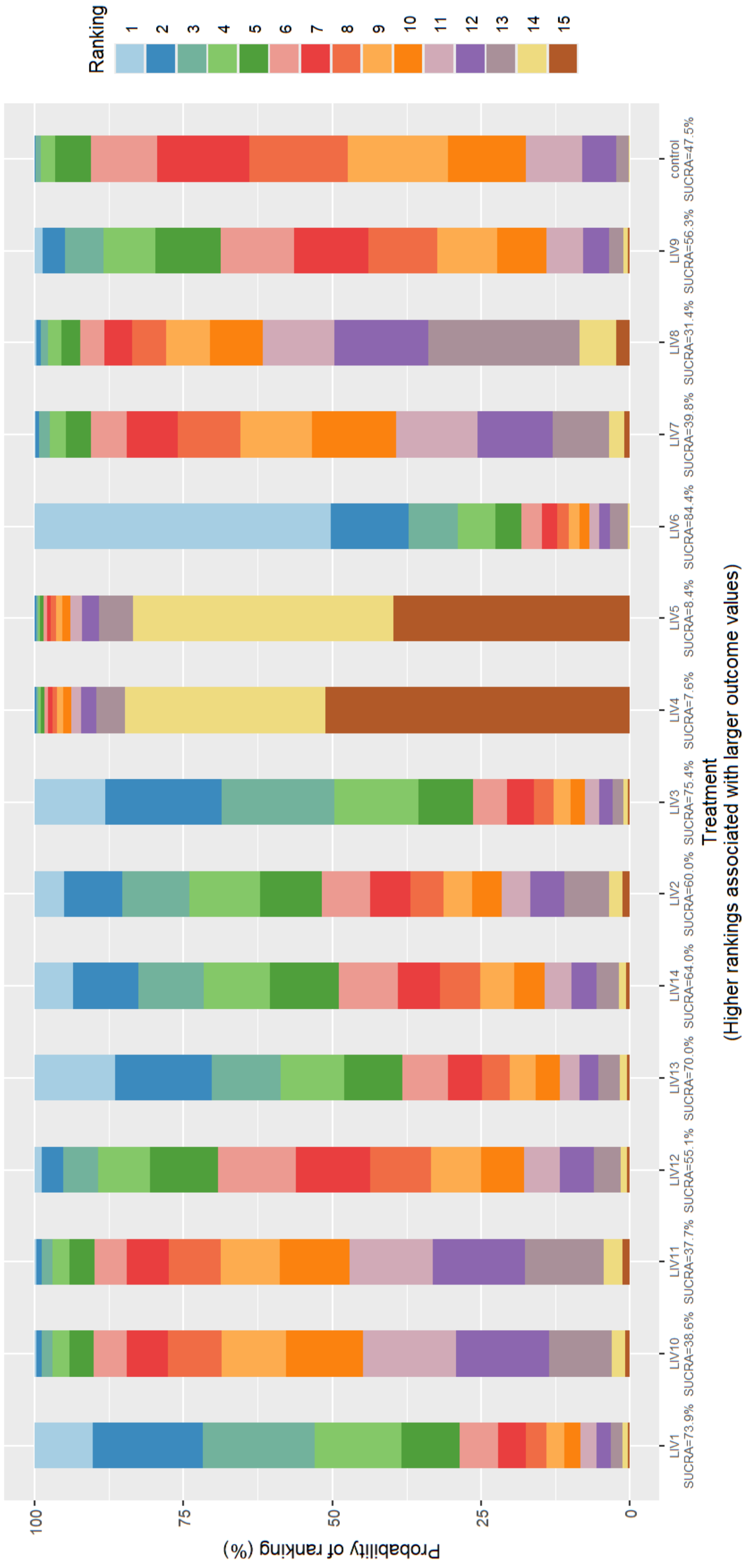

**Supplementary Figure 5. Heatmap for league table.** Mean differences with 95% credible interval for investigated LIVs in femurs **

** means *p* value < 0.01. The intensity of the color reflects the magnitude of the difference (darker = larger difference).

| Comparator \ Treatment | LIV6 | LIV3 | LIV1 | LIV13 | LIV14 | LIV2 | LIV9 | LIV12 | control | LIV7 | LIV10 | LIV11 | LIV8 | LIV5 | LIV4 |
|---|---|---|---|---|---|---|---|---|---|---|---|---|---|---|---|
| LIV6 | | -1.42 (-7.58, 4.87) | -1.50 (-7.58, 4.77) | -1.71 (-7.97, 4.79) | -2.21 (-8.29, 4.07) | -2.28 (-8.63, 4.11) | -2.74 (-8.25, 3.16) | -2.74 (-8.28, 2.95) | -3.08 (-7.96, 2.02) | -3.52 (-9.13, 2.17) | -3.54 (-8.99, 2.39) | -3.62 (-9.15, 2.27) | -3.99 (-9.57, 2.00) | **-7.06** (-11.82, -2.21) | **-7.36** (-12.41, -2.18) |
| LIV3 | 1.42 (-4.87, 7.58) | | -0.09 (-3.77, 3.62) | -0.32 (-5.73, 5.21) | -0.79 (-5.97, 4.42) | -0.91 (-4.91, 3.17) | -1.34 (-5.87, 3.45) | -1.31 (-6.04, 3.20) | -1.68 (-5.33, 2.07) | -2.05 (-6.84, 2.43) | -2.16 (-6.61, 2.67) | -2.21 (-6.85, 2.54) | -2.59 (-7.24, 2.38) | -5.64 (-11.44, 0.01) | -5.96 (-12.03, 0.03) |
| LIV1 | 1.50 (-4.77, 7.58) | 0.09 (-3.62, 3.77) | | -0.23 (-5.70, 5.23) | -0.69 (-5.89, 4.49) | -0.82 (-4.72, 3.22) | -1.24 (-5.80, 3.54) | -1.21 (-5.93, 3.28) | -1.60 (-5.24, 2.18) | -1.95 (-6.75, 2.41) | -2.06 (-6.53, 2.74) | -2.13 (-6.76, 2.57) | -2.51 (-7.15, 2.29) | -5.57 (-11.30, 0.13) | -5.84 (-11.86, 0.14) |
| LIV13 | 1.71 (-4.79, 7.97) | 0.32 (-5.21, 5.73) | 0.23 (-5.23, 5.70) | | -0.46 (-5.10, 4.15) | -0.59 (-6.28, 5.10) | -1.01 (-5.04, 3.25) | -1.02 (-5.50, 3.25) | -1.37 (-5.34, 2.66) | -1.74 (-6.09, 2.23) | -1.81 (-5.83, 2.38) | -1.87 (-6.05, 2.32) | -2.24 (-6.57, 2.17) | -5.35 (-11.30, 0.49) | -5.70 (-11.78, 0.56) |
| LIV14 | 2.21 (-4.07, 8.29) | 0.79 (-4.42, 5.97) | 0.69 (-4.49, 5.89) | 0.46 (-4.15, 5.10) | | -0.12 (-5.54, 5.33) | -0.53 (-4.24, 3.38) | -0.54 (-4.66, 3.44) | -0.89 (-4.52, 2.83) | -1.28 (-5.14, 2.30) | -1.35 (-4.99, 2.49) | -1.41 (-5.23, 2.45) | -1.77 (-5.73, 2.30) | -4.89 (-10.63, 0.82) | -5.16 (-11.17, 0.81) |
| LIV2 | 2.28 (-4.11, 8.63) | 0.91 (-3.17, 4.91) | 0.82 (-3.22, 4.72) | 0.59 (-5.10, 6.28) | 0.12 (-5.33, 5.54) | | -0.43 (-5.24, 4.57) | -0.44 (-5.28, 4.27) | -0.78 (-4.73, 3.27) | -1.17 (-6.10, 3.44) | -1.24 (-5.92, 3.74) | -1.30 (-6.10, 3.65) | -1.67 (-6.48, 3.39) | -4.78 (-10.78, 1.10) | -5.04 (-11.35, 1.13) |
| LIV9 | 2.74 (-3.16, 8.25) | 1.34 (-3.45, 5.87) | 1.24 (-3.54, 5.80) | 1.01 (-3.25, 5.04) | 0.53 (-3.38, 4.24) | 0.43 (-4.57, 5.24) | | 0.01 (-3.62, 3.18) | -0.36 (-3.23, 2.36) | -0.73 (-3.87, 1.80) | -0.81 (-3.63, 2.04) | -0.88 (-3.91, 1.96) | -1.24 (-4.41, 1.86) | -4.34 (-9.73, 0.68) | -4.65 (-10.23, 0.80) |
| LIV12 | 2.74 (-2.95, 8.28) | 1.31 (-3.20, 6.04) | 1.21 (-3.28, 5.93) | 1.02 (-3.25, 5.50) | 0.54 (-3.44, 4.66) | 0.44 (-4.27, 5.28) | -0.01 (-3.18, 3.62) | | -0.38 (-2.92, 2.44) | -0.74 (-4.15, 2.46) | -0.84 (-3.89, 2.68) | -0.89 (-4.16, 2.66) | -1.27 (-4.67, 2.50) | -4.32 (-9.43, 0.81) | -4.62 (-10.01, 0.92) |
| control | 3.08 (-2.02, 7.96) | 1.68 (-2.07, 5.33) | 1.60 (-2.18, 5.24) | 1.37 (-2.66, 5.34) | 0.89 (-2.83, 4.52) | 0.78 (-3.27, 4.73) | 0.36 (-2.36, 3.23) | 0.38 (-2.44, 2.92) | | -0.38 (-3.32, 2.12) | -0.46 (-3.04, 2.26) | -0.53 (-3.28, 2.27) | -0.89 (-3.84, 2.12) | -3.98 (-8.47, 0.31) | -4.28 (-9.15, 0.42) |
| LIV7 | 3.52 (-2.17, 9.13) | 2.05 (-2.43, 6.84) | 1.95 (-2.41, 6.75) | 1.74 (-2.23, 6.09) | 1.28 (-2.30, 5.14) | 1.17 (-3.44, 6.10) | 0.73 (-1.80, 3.87) | 0.74 (-2.46, 4.15) | 0.38 (-2.12, 3.32) | | -0.10 (-2.53, 2.99) | -0.14 (-2.79, 2.86) | -0.51 (-3.26, 2.80) | -3.59 (-8.60, 1.63) | -3.87 (-9.15, 1.78) |
| LIV10 | 3.54 (-2.39, 8.99) | 2.16 (-2.67, 6.61) | 2.06 (-2.74, 6.53) | 1.81 (-2.38, 5.83) | 1.35 (-2.49, 4.99) | 1.24 (-3.74, 5.92) | 0.81 (-2.04, 3.63) | 0.84 (-2.68, 3.89) | 0.46 (-2.26, 3.04) | 0.10 (-2.99, 2.53) | | -0.06 (-2.99, 2.67) | -0.43 (-3.47, 2.51) | -3.53 (-8.85, 1.47) | -3.84 (-9.40, 1.63) |
| LIV11 | 3.62 (-2.27, 9.15) | 2.21 (-2.54, 6.85) | 2.13 (-2.57, 6.76) | 1.87 (-2.32, 6.05) | 1.41 (-2.45, 5.23) | 1.30 (-3.65, 6.10) | 0.88 (-1.96, 3.91) | 0.89 (-2.66, 4.16) | 0.53 (-2.27, 3.28) | 0.14 (-2.86, 2.79) | 0.06 (-2.67, 2.99) | | -0.38 (-3.40, 2.80) | -3.47 (-8.70, 1.64) | -3.75 (-9.27, 1.76) |
| LIV8 | 3.99 (-2.00, 9.57) | 2.59 (-2.38, 7.24) | 2.51 (-2.29, 7.15) | 2.24 (-2.17, 6.57) | 1.77 (-2.30, 5.73) | 1.67 (-3.39, 6.48) | 1.24 (-1.86, 4.41) | 1.27 (-2.50, 4.67) | 0.89 (-2.12, 3.84) | 0.51 (-2.80, 3.26) | 0.43 (-2.51, 3.47) | 0.38 (-2.80, 3.40) | | -3.09 (-8.55, 2.10) | -3.40 (-9.13, 2.14) |
| LIV5 | **7.06** (2.21, 11.82) | 5.64 (-0.01, 11.44) | 5.57 (-0.13, 11.30) | 5.35 (-0.49, 11.30) | 4.89 (-0.82, 10.63) | 4.78 (-1.10, 10.78) | 4.34 (-0.68, 9.73) | 4.32 (-0.81, 9.43) | 3.98 (-0.31, 8.47) | 3.59 (-1.63, 8.60) | 3.53 (-1.47, 8.85) | 3.47 (-1.64, 8.70) | 3.09 (-2.10, 8.55) | | -0.30 (-4.78, 4.23) |
| LIV4 | **7.36** (2.18, 12.41) | 5.96 (-0.03, 12.03) | 5.84 (-0.14, 11.86) | 5.70 (-0.56, 11.78) | 5.16 (-0.81, 11.17) | 5.04 (-1.13, 11.35) | 4.65 (-0.80, 10.23) | 4.62 (-0.92, 10.01) | 4.28 (-0.42, 9.15) | 3.87 (-1.78, 9.15) | 3.84 (-1.63, 9.40) | 3.75 (-1.76, 9.27) | 3.40 (-2.14, 9.13) | 0.30 (-4.23, 4.78) | |

**Supplementary Figure 6. Heatmap for league table.** Mean differences with 95% credible interval for investigated LIVs in tibias ** ** means *p* value < 0.01. The intensity of the color reflects the magnitude of the difference (darker = larger difference).

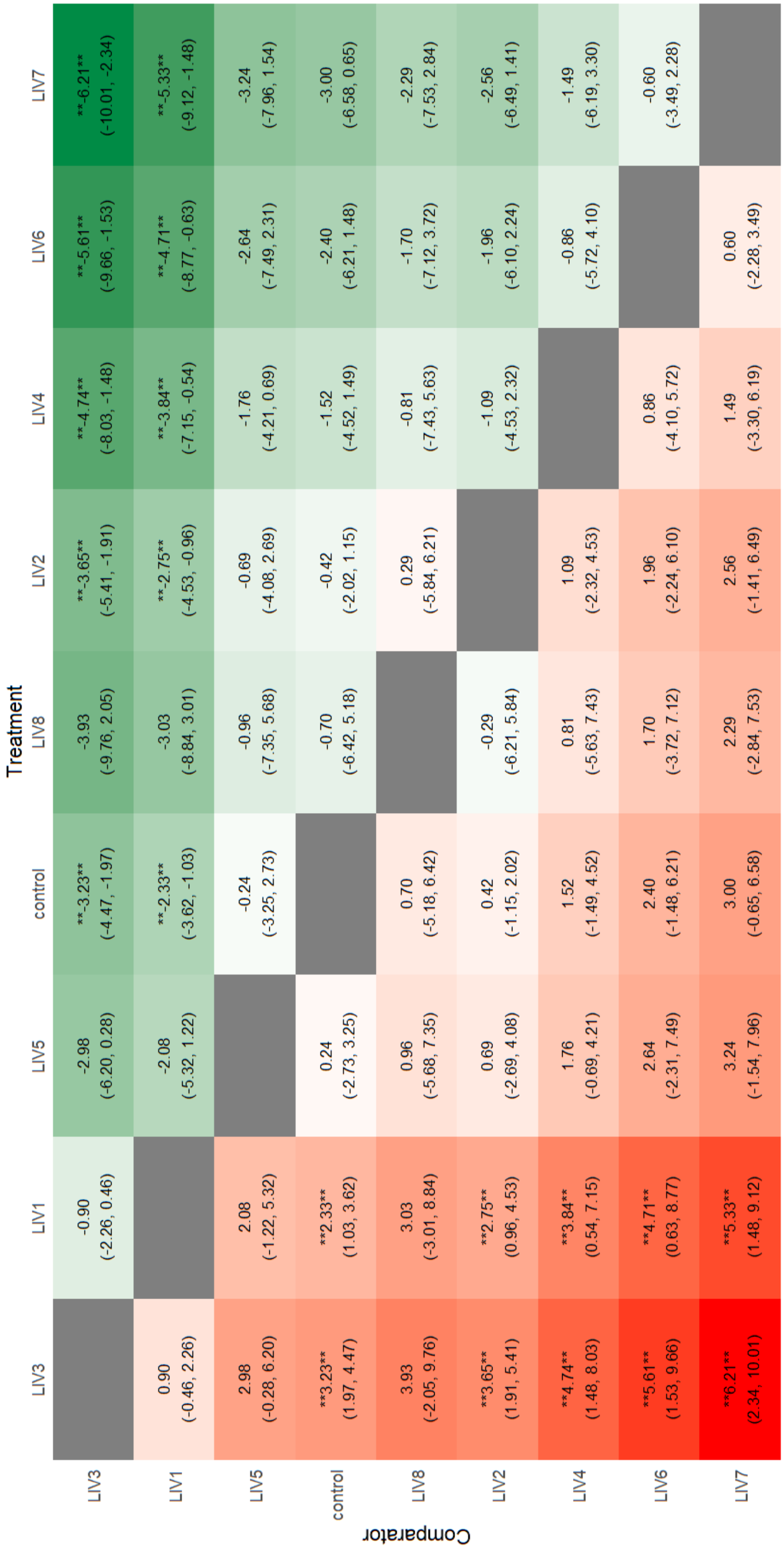

**Supplementary Table 1A:** Gelman-Rubin and Geweke results

| | Point estimations for Gelman-Rubin results | Geweke statistics Chain 1 | Geweke statistics Chain 1 | Geweke statistics Chain 1 |
|---|---|---|---|---|
| d[2] | 1.00021 | -0,26355 | 1,02830 | 1,06961 |
| d[3] | 1.00033 | -0,44477 | -0,08478 | 0,34419 |
| d[4] | 1.00053 | -0,72225 | 0,71815 | 1,83287 |
| d[5] | 1.00022 | -0,24916 | 0,53583 | -0,83289 |
| d[6] | 1.00019 | -0,84735 | 0,23117 | -1,22103 |
| d[7] | 1.00011 | 0,30059 | -0,40111 | 0,32860 |
| d[8] | 1.00015 | 0,27099 | -0,27563 | 0,37565 |
| d[9] | 1.00031 | 0,58747 | -0,16008 | -0,01950 |

**Supplementary Table 1B:** Gelman-Rubin and Geweke results

| | Point estimations for Gelman-Rubin results | Geweke statistics Chain 1 | Geweke statistics Chain 1 | Geweke statistics Chain 1 |
|---|---|---|---|---|
| d[2] | 1.00028 | -0.81172 | 2.15337 | -0.85434 |
| d[3] | 1.00169 | 0.33455 | -0.72021 | -0.22409 |
| d[4] | 1.00099 | 0.78693 | -0.01382 | 0.33868 |
| d[5] | 1.00100 | 3.75084 | -0.72440 | -1.12956 |
| d[6] | 1.00051 | 0.78684 | 1.30815 | 0.17349 |
| d[7] | 1.00034 | 1.36512 | 1.21298 | -1.20255 |
| d[8] | 1.00049 | -0.94934 | 1.90170 | -1.20371 |
| d[9] | 1.00070 | -0.76054 | 2.29222 | -0.64559 |
| d[10] | 1.00091 | 1.03317 | -1.48183 | -1.87766 |
| d[11] | 1.00016 | 0.27690 | -1.30609 | -0.67296 |

| | | | | |
|---|---|---|---|---|
| d[12] | 1.00045 | -0.21912 | -1.00288 | -0.32312 |
| d[13] | 1.00033 | 0.77443 | 0.14591 | -0.52226 |
| d[14] | 1.00099 | 0.20935 | -1.31724 | 0.86011 |
| d[15] | 1.00079 | 0.15637 | 0.35475 | -0.25669 |
| sigma | 1.00076 | -0.50840 | -1.49509 | 1.17367 |
